# Defining the roles of the Integrator, NEXT, and nuclear exosome complexes in *Drosophila* oogenesis

**DOI:** 10.1101/2025.03.05.641556

**Authors:** Yongjin Lee, Adriano Biasini, Cindy Tipping, Seong Hyeon Hong, Phillip D. Zamore

**Affiliations:** RNA Therapeutics Institute, University of Massachusetts Chan Medical School, 368 Plantation Street, Worcester, MA 01605, USA; Howard Hughes Medical Institute, University of Massachusetts Chan Medical School, 368 Plantation Street, Worcester, MA 01605, USA

**Keywords:** Integrator, NEXT, Exosome, Ars2, IntS11, Rrp6, Dis3, cryptic transcript, read-through transcription, Oocyte localization, RNA 3′ end processing, small nuclear RNA

## Abstract

Nuclear RNA homeostasis depends on the balance of transcription, RNA processing, degradation, and transport between the nucleus and cytoplasm. RNA degradation directed by the Integrator, nuclear exosome targeting (NEXT), and nuclear exosome complexes controls the accumulation of aberrant nuclear RNA. Here, we report that *Drosophila* oogenesis requires the Integrator, NEXT, and nuclear exosome complexes. Depletion of Integrator, NEXT, or nuclear exosome components in *Drosophila* female germ cells causes infertility and accumulation of 3′ extended snRNAs, promoter upstream transcripts (PROMPTs), and cryptic transcripts. Our data highlight the essential role of nuclear RNA degradation and processing in *Drosophila* oogenesis and provide a catalog of RNAs whose nuclear levels are regulated by these three complexes. We propose that Integrator, NEXT, and the nuclear exosome support oogenesis by ensuring that inappropriate transcription does not overwhelm the limited supply of proteins that bind, process, and traffic RNA.

## INTRODUCTION

The steady-state level of nuclear RNA reflects the combined rates of transcription, RNA processing, intranuclear degradation, and export to the cytoplasm. Many classes of RNA can only exit the nucleus after they have had intervening sequences removed and 3′ ends appropriately defined.(Lei and Silver, 2002; Hocine et al., 2010; Köhler and Hurt, 2007; Millevoi and Vagner, 2010) In animals, RNA polymerase II (Pol II) produces primary transcripts that extend beyond the 3′ boundary of the functional RNA species; endonucleolytic cleavage or exonucleolytic trimming serves to convert these primary transcripts into mature, functional RNAs. Moreover, Pol II generates short-lived RNAs whose degradation is thought to be essential for nuclear RNA homeostasis. These include promoter upstream transcripts (PROMPTs) originating from bidirectional promoters and cryptic transcripts of little or no function. Three multi-protein ribonuclease complexes play essential roles in nuclear RNA homeostasis: the Integrator complex, the nuclear exosome targeting (NEXT) complex, and the nuclear exosome.

Integrator plays diverse roles in transcription and RNA processing, including reducing the effective rate of transcription by promoting premature termination of individual genes under specific environmental conditions and eliminating ‘downstream of gene’ (DoG) read-through transcripts.(Lykke-Andersen et al., 2021; Dasilva et al., 2021; Rosa-Mercado et al., 2021; Beckedorff et al., 2020; Tatomer et al., 2019; Elrod et al., 2019) Furthermore, Integrator regulates transcription of telomerase RNA and replication-dependent histone mRNA and participates in enhancer RNA biogenesis and transcript isoform switching.(Barra et al., 2020; Rubtsova et al., 2019; Skaar et al., 2015; Lai et al., 2015) A core function of Integrator is establishing the 3′ ends of the spliceosomal small nuclear RNAs (snRNAs) synthesized by RNA Pol II.(Baillat et al., 2005) Spliceosomal snRNAs are transcribed with 3′ extensions that are removed by Integrator, and depletion of Integrator subunits leads to accumulation of 3′ extended snRNAs in human and *Drosophila melanogaster* cell lines, and in vivo in *Caenorhabditis elegans* and the planarian *Schmidtea mediterranea*.(Pfleiderer and Galej, 2021; Gómez-Orte et al., 2019; Schmidt et al., 2018; O’Reilly et al., 2014; Baillat et al., 2005)

Integrator comprises fifteen subunits (INTS1–INTS15) that are organized into multiple modules.(Fianu et al., 2024; Zheng et al., 2023; Azuma et al., 2023; Offley et al., 2023; Baillat et al., 2005; Chen et al., 2012) INTS1, INTS2 and INTS5–INTS8 form the core of Integrator. The core proteins INTS2, INTS5, INTS6, and INTS8 together bind the Protein Phosphatase 2A (PP2A) subunits PP2A-A and PP2A-C; Integrator regulates transcriptional initiation, pausing, and elongation by directing PP2A to dephosphorylate Pol II.(Fianu et al., 2024; Zheng et al., 2023; Fianu et al., 2021; Zheng et al., 2020) INTS10, INTS13, INTS14 and INTS15 comprise the tail module, which bridges the core and catalytic modules.(Fianu et al., 2024; Sabath et al., 2020) INTS4, INTS9, and the endoribonuclease INTS11 compose the Integrator catalytic module.(Pfleiderer and Galej, 2021; Albrecht et al., 2018; Wu et al., 2017; Albrecht and Wagner, 2012) INTS9 and INTS11 are structurally homologous to the cleavage and polyadenylation specificity factor components CPSF100 and CPSF73, which together define mRNA 3′ ends and promote polyadenylation.(Albrecht and Wagner, 2012; Dominski et al., 2005; Baillat et al., 2005) Unlike CPSF, which recognizes the polyadenylation signal on mRNA, how the Integrator catalytic module recognizes its substrates remains unknown.

The assembly factors BRAT1 (BRCA1-associated ATM activator 1) and WDR73 (WD repeat domain 73) facilitate nuclear import of the INTS9•INTS11 dimer. BRAT1 binding also blocks the active site of the INTS11 ribonuclease, preventing INTS11 from degrading cytoplasmic RNA.(Sabath et al., 2024) In the nucleus, BRAT1 dissociates and INTS4 joins INTS9•INTS11 to generate the trimeric catalytic module, which can then join the Integrator core module.(Dokaneheifard et al., 2024; Lin et al., 2024; Sabath et al., 2024; Cihlarova et al., 2022; Tilley et al., 2021; Pfleiderer and Galej, 2021)

A major function of Integrator is to reduce gene expression by prematurely terminating transcription. Productive transcription requires the conversion of paused Pol II into actively elongating polymerase, a process that requires phosphorylation of the Pol II carboxy-terminal domain (CTD). CTD phosphorylation allows binding of the elongation factors SPT6 (Suppressor of Ty6) and PAF1 (Polymerase-Associated Factor 1).(Vos et al., 2018; Sun et al., 2010) The Integrator complex binds the serine/threonine protein phosphatase PP2A, positioning it to dephosphorylate the CTD of Pol II, thereby blocking binding of SPT6 and PAF1 and terminating transcription.(Fianu et al., 2024; Fianu et al., 2021; Huang et al., 2020; Zheng et al., 2020; Cortazar et al., 2019; Yamamoto et al., 2014; Egloff et al., 2012; Egloff et al., 2010)

The NEXT complex acts as an adapter to link non-polyadenylated RNA substrates to the nuclear exosome. NEXT comprises the RNA helicase MTR4, the RNA-binding protein RBM7, and the zinc-finger protein ZCCHC8.(Wu et al., 2020; Meola et al., 2016; Lubas et al., 2011) ZCCHC8 binds both MTR4 and RBM7, positioning RBM7 next to the MTR4 helicase domain, which may facilitate RNA substrate recognition.(Gerlach et al., 2022; Puno and Lima, 2022; Puno and Lima, 2018; Falk et al., 2016; Tiedje et al., 2015) The NEXT complex binds the 3′ ends of RNAs containing poly(U) rather than poly(A) stretches and directs the substrate to the nuclear exosome for degradation.(Hrossova et al., 2015) The RNA-binding protein Ars2 (arsenic-resistance protein 2) and the zinc-finger protein ZC3H18, which copurify with cap-binding protein 20 (CBP20), cap-binding protein 80 (CBP80), Mtr4, and RBM7, together connect RNA substrates to the NEXT complex by binding both the nuclear RNA cap-binding complex (CBC) and NEXT.(Andersen et al., 2013) Ars2 is required for cell proliferation, early mammalian development, viral miRNA/siRNA biogenesis, and 3’ end formation of replication-dependent histone pre-mRNA.(Gruber et al., 2012; Sabin et al., 2009; Gruber et al., 2009; Wilson et al., 2008) In *Arabidopsis*, the Ars2 ortholog, SERRATE regulates miRNA biogenesis by linking primary miRNA precursors to the NEXT complex.(Fang and Spector, 2007)

The nuclear RNA exosome, a conserved multi-protein ribonuclease complex, regulates the accumulation of nuclear RNA by degrading cryptic or defective transcripts.(Kilchert et al., 2016) Recruitment of specific RNAs or RNA classes to the exosome requires adapter complexes that bind both the exosome and the RNA substrate.(Wu et al., 2020) For example, the NEXT complex binds PROMPTs (promoter upstream transcripts), short RNAs generated by divergent transcription of promoters driving mRNA synthesis, and directs them to the nuclear exosome for destruction. The two nuclear exosome RNases, Rrp6 and Dis3, have distinct substrate specificities. A 3′-to-5′ exoribonuclease, Rrp6 preferentially degrades 3′ hydroxyl-bearing RNAs, while Dis3 can act as both an endonuclease and a 3′-to-5′ exoribonuclease, targeting both 3′ hydroxy and 3′ phosphorylated RNA.(Zinder et al., 2016; Wasmuth et al., 2014) Dis3 degrades prematurely terminated polyadenylated RNAs and RNA produced by divergent transcription at the promoters of protein coding genes.(Sigova et al., 2013; Chiu et al., 2018) In mammals, maturation of 5.8S ribosomal RNA (rRNA) and small nucleolar RNAs require Rrp6 (EXOSC10).(Davidson et al., 2019)

Here, we define the subunit compositions of the Integrator, NEXT, and exosome complexes in the fly ovary. In addition to identifying known components of each complex, our data identify new interactions both among the complexes and with proteins previously not linked to nuclear RNA regulation. In vivo, Integrator, NEXT, and the nuclear exosome are required in the germ line for oogenesis in *D. melanogaster*. Female flies depleted in the germ line of individual components of these complexes are sub-fertile or sterile, and generally lay few eggs, which rarely hatch; none develop to adulthood. Although the oocyte is thought to be transcriptionally quiescent for most genes, Ars2, ZC3H18, Rrp6, and Dis3—proteins that act on newly transcribed RNAs— accumulate in the oocyte nucleus. We define the global consequences for RNA homeostasis—including defective snRNA 3′ end processing and accumulation of cryptic transcripts, RNA from promoters and introns—upon germ-line depletion of individual Integrator, NEXT, or nuclear exosome components. Our data provides a resource for future investigations of the interactions and functions of the Integrator, NEXT, and nuclear exosome complexes as well as the contributions of nuclear RNA homeostasis to *Drosophila* female germ-line development.

## RESULTS

### Fly Inst11 and Ars2 copurify with known components of the Integrator and NEXT complexes

To test whether the composition of the Integrator, NEXT, and nuclear exosome complexes in *Drosophila* ovaries mirrors that determined in fly and human cultured cells, we developed a novel copurification/mass spectrometry strategy. We generated fly lines expressing Twin-Strep-3XFLAG-tagged IntS11, Ars2, Rrp6, or Dis3 (Figure 1A) and treated their ovaries with the homobifunctional crosslinker dithio-bis-maleimidoethane (DTME). We note that all five knock-in, epitope tagged fly lines were homozygous viable and fully fertile. Next, we purified each epitope-tagged protein using Strep-Tactin beads in the presence of 0.5% sodium dodecyl sulfate (SDS). As a control, we performed the crosslinking and Strep-Tactin purification using ovaries from *w^1118^*flies, which express no epitope-tagged proteins. We used LC-MS/MS mass spectrometry to identify copurifying proteins (Figure 1B). The use of SDS eliminated the high non-specific background typically associated with co-immunoprecipitation/mass spectrometry experiments, allowing us to consider only proteins copurifying with an epitope-tagged protein but not detectably recovered in the non-tagged control.

**Figure 1.**
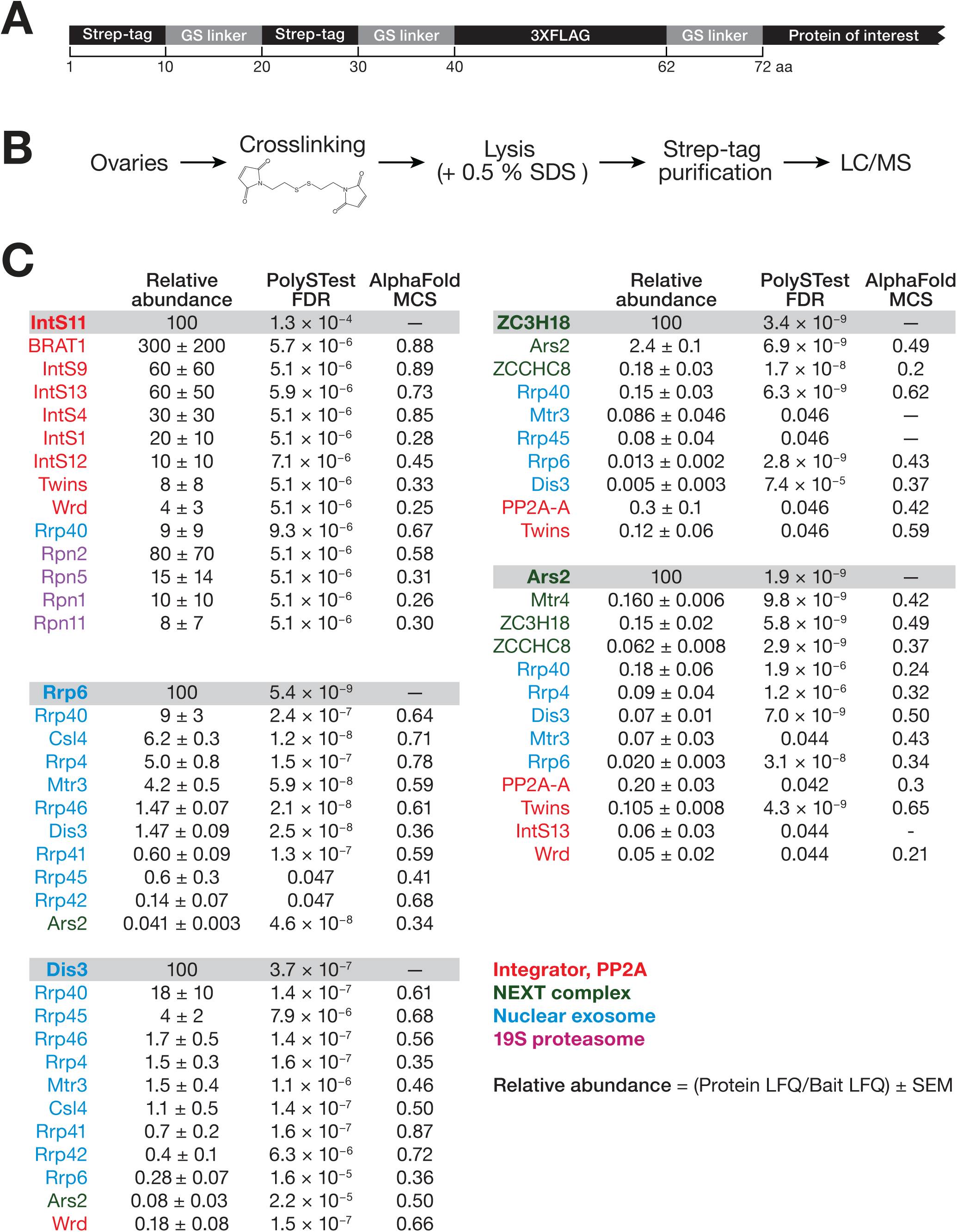
Proteins that interact with Integrator, NEXT, or the nuclear exosome. (A) Epitope-tagging strategy used to generate knock-in fly strains. (B) Strategy used to identify copurifying proteins. (C) Proteins that copurified with epitope-tagged target proteins (FDR < 0.05). The Twin-Strep-3XFLAG-tagged protein is listed in bold at the top of each section. Protein abundance was calculated by label-free quantification (LFQ) and reported as mean ± SEM relative to the abundance of the tagged protein. Significance testing was performed using PolySTest to obtain false discovery rate (FDR).(Schwämmle et al., 2020) AlphaFold-multimer predicts the surface interface probability between target and copurified proteins. The highest modeling confidence scores (MCS) obtained from the analysis of multiple seeds are reported. Red, Integrator and PP2A subunits; dark green, NEXT components; blue, exosome components; purple, 19S proteasome regulatory particle non-ATPases.

Five of the 14 known subunits (IntS1, IntS4, IntS9, IntS12, and IntS13) of the Integrator complex copurified with IntS11 in vivo (Figure 1C and Table S1); the cleavage module components IntS4 and IntS9 were among the most abundant copurifying proteins, corroborating the previous finding that IntS4 binds the IntS11•IntS9 heterodimer to form a stable complex.(Fianu et al., 2024; Albrecht et al., 2018) BRAT1, a protein that facilitates nuclear import of the IntS11•IntS9 heterodimer, copurified at comparable abundance to Ints4 and Ints9.(Sabath et al., 2024) The Integrator complex binds the serine/threonine protein phosphatase PP2A(Fianu et al., 2024; Fianu et al., 2021; Huang et al., 2020); two PP2A regulatory subunits copurified with IntS11: Twins (tws), the fly ortholog of human PP2A-B (B55), and Well rounded (wrd), the ortholog of PP2A-B′ (B56).(Cho and Xu, 2007) The nuclear exosome subunit Rrp40 also copurified with IntS11, consistent with the observation that Integrator binds the nuclear RNA exosome.(Tatomer et al., 2019)

IntS11 also copurified with eight non-ATPase components of the 19S proteasome regulatory complex: Rpn1, Rpn2, Rpn3, Rpn5, Rpn8, Rpn10, Rpn11, and Rpn12. None were detected in the non-tagged ovary control. During *Drosophila* oogenesis, subunits of the 19S proteasome localize to the cytoplasm of the germ-line nurse cells and the somatic follicular epithelium, while the complete 26S proteasome accumulates in the oocyte nucleus, peaking at stage 7, after which the number of oocytes with detectable nuclear 26S proteasome proteins declines.(Arkinson et al., 2025; Uriarte et al., 2021; Bard et al., 2018; Livneh et al., 2016; Reits et al., 1997; Adám et al., 2004) The 26S proteasome also accumulates in the nuclei of the columnar follicular cells that surround the oocyte. Although the Integrator complex was discovered because of its association with the 19S proteasome protein DSS1 (Deleted in split hand/split foot 1)(Baillat et al., 2005), we did not detect the *Drosophila* ortholog of DSS1, Sem1, among the IntS11-copurifying proteins. The failure to recover Sem1 with IntS11 may reflect an absence of suitable cysteine residues to facilitate its crosslinking.

We used AlphaFold Multimer(Evans et al., 2021; Jumper et al., 2021) to test whether copurification of 19S proteasome proteins might plausibly reflect direct interactions with IntS11 (Figure 1C and Table S2). Rpn1, Rpn2, Rpn3, Rpn5, Rpn8, Rpn10, Rpn11, and Rpn12 were each co-folded with IntS11 to obtain a model confidence score (MCS).(Evans et al., 2021) The modeling-based analysis predicted that four of the eight copurifying proteasome regulatory subunits directly bind IntS11 with MCS >0.5 (Rpn2; MCS = 0.58, Rpn3; MCS = 0.67, Rpn8; MCS = 0.51, and Rpn10; MCS = 0.7). Although Rpn7 did not copurify with IntS11, AlphaFold Multimer also predicts that Rpn7 directly binds IntS11 (MCS = 0.62).

The NEXT complex, which comprises Mtr4, ZCCHC8, and RBM7, requires Ars2 to bind to its RNA substrates.(Andersen et al., 2013) The Drosophila RNAi Screening Center Integrative Ortholog Prediction Tool (DIOPT) predicts that the CG1677 C3H1 zinc finger sequence shares 30% identity with human ZC3H18 (overall identity, 24%; similarity, 34%).(Hu et al., 2011) Twin-Strep-tagged Ars2 and CG1677 reciprocally copurified, supporting the view that CG1677 is the *Drosophila* ortholog of the human Ars2-binding protein ZC3H18 (Figure 1C and Table S2). Henceforth, we refer to CG1677 as ZC3H18. The NEXT complex component ZCCHC8 copurified with both Ars2 and ZC3H18, and Mtr4 copurified with Ars2. Five of the 10 components of the nuclear exosome (Rrp4, Rrp6, Rrp40, Dis3 and Mtr3) copurified with Ars2. Similarly, the nuclear exosome proteins, Rrp6, Rrp40, Rrp45, Dis3 and Mtr3 copurified with ZC3H18. Our data suggest that, as in mammals, fly ZC3H18 and Ars2 link the NEXT complex to the nuclear exosome. CG11454, the *Drosophila* ortholog of the mammalian NEXT component RBM7, did not detectably copurify with either Ars2 or ZC3H18. This may reflect a difference in the composition of the human and fly NEXT complexes or simply a lack of accessible cysteines for crosslinking by DTME.

The nuclear cap-binding complex (CBC) component NCBP1 also copurified with Ars2 and ZC3H18. CBC binds the newly synthesized 5′ 7-methyl guanosine cap characteristic of Pol II transcripts and participates in many aspects of nuclear RNA processing, including pre-mRNA splicing, mRNA 3′ end processing, microRNA biogenesis, and RNA export.(Dubiez et al., 2024; Hallais et al., 2013; Gruber et al., 2009) Binding of NCBP1 to Ars2 or ZC3H18 is unlikely to be mediated by RNA, because our lysates were incubated with RNases A and T1 and our purification conditions are designed to require DTME crosslinking via sulfhydryl groups. In mammals, the CBC binds Ars2, which, together with ZC3H18, links it to the exosome.(Polák et al., 2023; Winczura et al., 2018; Schulze and Cusack, 2017; Andersen et al., 2013; Hallais et al., 2013)

All known nuclear exosome proteins—Mtr3, Csl4, Dis3, Rrp4, Rrp6, Rrp40, Rrp41, Rrp42, Rrp45, and Rrp46—copurified with the nuclear exosome nucleases Rrp6 and Dis3. Moreover, Mtr3, Rrp40, Rrp6, and Dis3 copurified with Ars2 and ZC3H18, consistent with the NEXT complex targeting RNA for degradation by delivering it to the nuclear exosome.(Andersen et al., 2013; Lubas et al., 2011)

### Integrator, NEXT, and nuclear exosome components are associated with proteins involved in oogenesis and RNA processing

Hundreds of proteins copurified (FDR < 0.01) with the epitope-tagged proteins and were either depleted from or not recovered at all in the non-tagged control: 154 for IntS11, 472 for Ars2, 379 for ZC3H18, 286 for Rrp6, and 303 for Dis3. More than half the proteins copurifying with IntS11 also copurified with other epitope-tagged proteins: 79% with Ars2, 79% with ZC3H18, 70% with Rrp6, and 67% with Dis3 (Figure 2A). Similarly, proteins that copurified with Ars2 often copurified with IntS11 (79%), Rrp6 (85%), Dis3 (88%), and ZC3H18 (80%). Moreover, 74% of proteins that copurified with Rrp6 also copurified with Dis3. Proteins that copurified with Integrator, NEXT, and nuclear exosome components participate in oogenesis, mRNA and protein transport, RNA processing, transcription regulation, and splicing (Figure 2B). For example, Rumpelstiltskin (Rump), Cup, and Bicaudal (Bic), which all copurified with Rrp6, Dis3, ZC3H18, and Ars2, regulate the localization and translation of *oskar* (*osk*) mRNA, which is essential for embryonic patterning and germ cell determination.(Saffman et al., 1998) Hrb27C and Squid (sqd), which copurified with all three complexes, function in the localization and translation of *gurken* (*grk*) mRNA. Grk establishes the dorsal-ventral axis during fly oogenesis (Figure 2B).(Goodrich et al., 2004; Neuman-Silberberg and Schupbach, 1994; Neuman-Silberberg and Schüpbach, 1993) Lost also copurified with all three complexes; it interacts with Rump and is involved in the localization of mRNAs, including *osk*, to the oocyte posterior.(Sinsimer et al., 2011) Rings lost (Rngo) also copurified with all three complexes; it is required for germ-line cyst formation and ring canal development during oogenesis.(Morawe et al., 2011)

**Figure 2.**
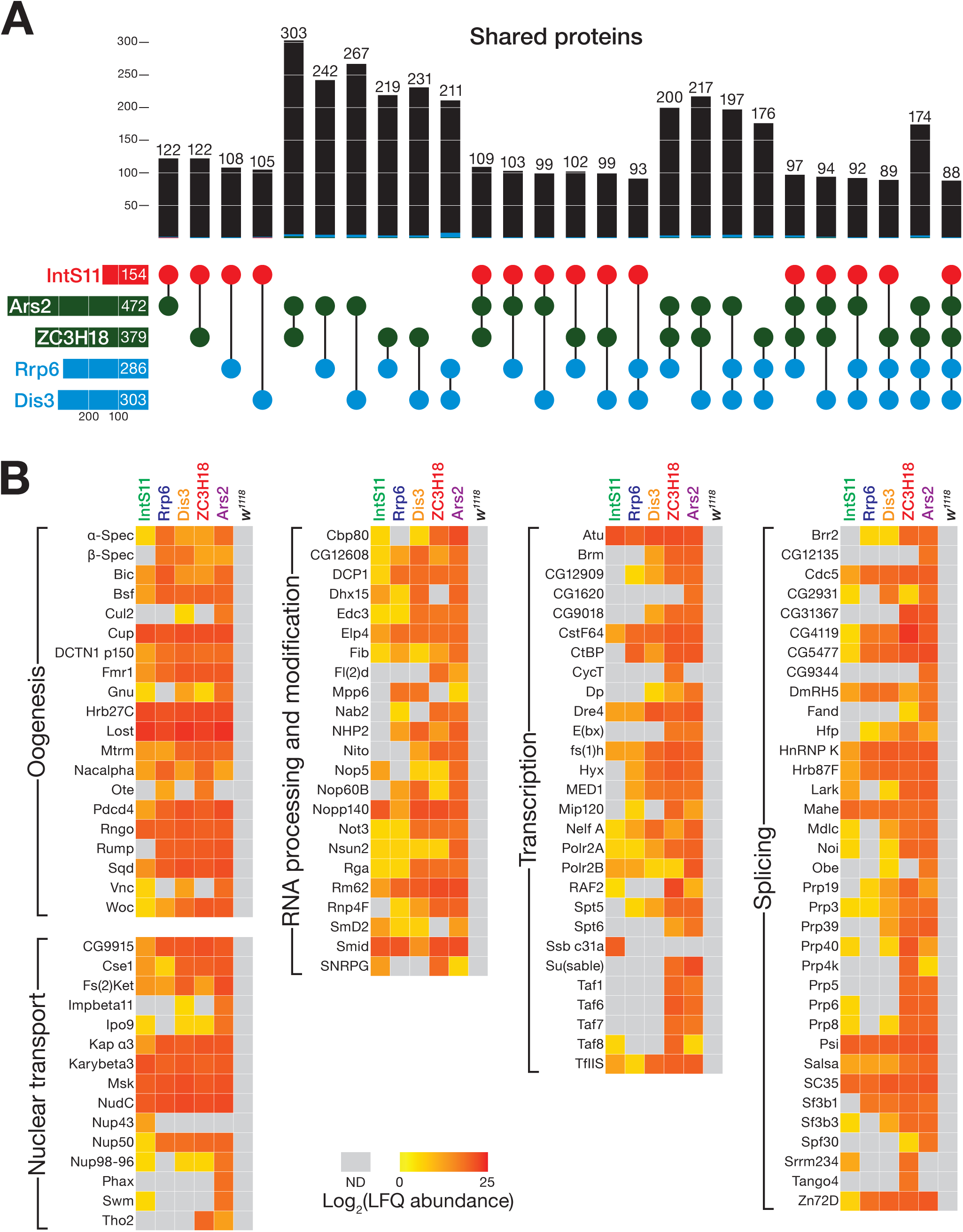
Integrator, NEXT, and nuclear exosome components bind to proteins functioning in oogenesis, transport, transcription, and RNA processing. (A) Upset plot(Lex et al., 2014) of the overlap among proteins copurifying with each tagged protein: IntS11 (red), NEXT (green); nuclear exosome (blue). Left, total proteins copurified with each tagged protein. Top, number of proteins copurifying with more than one tagged protein. (B) Copurified proteins were grouped by their functions using UniProt, FlyBase, and Gene Ontology annotations. Heatmap indicates the log_2_(relative abundance) of the proteins that copurified with each epitope-tagged protein. Gray, not detected.

Consistent with Integrator, NEXT, and the nuclear exosome acting co-transcriptionally, the Pol II subunits Polr2A and Polr2B, the transcription initiation factors Taf1, Taf6, Taf7, and Taf8, and the transcription elongation factors TfIIS, SPT5, and SPT6 copurified with components of all three complexes. A ring of seven small nuclear ribonucleoprotein Sm proteins bind the Sm site in the U1, U2, U4, and U5 snRNAs and participate in snRNA biogenesis and splicing.(Jurica and Moore, 2003; Pellizzoni et al., 2002) SmD2 copurified with IntS11, Rrp6, Dis3, and Ars2, and Small nuclear ribonucleoprotein G (SNRPG) copurified with IntS11, ZC3H18, and Ars2. The Sm ring protects mature snRNAs from 3′-to-5′ degradation by the nuclear exosome.(Coy et al., 2013) Sm proteins bind to snRNAs allowing them to assemble into small nuclear ribonucleoproteins (snRNPs), be re-imported into the nucleus, and localize together with other spliceosomal proteins to the Cajal body.(Matera and Wang, 2014) Spliceosomal proteins also copurified with Ars2 and ZC3HC18 (Figure 2B).

rRNA processing proteins also copurified with components of Integrator, NEXT, and the nuclear exosome. The nucleolar protein Nopp140 copurified with components of all three complexes: Nop5 copurified with IntS11, Dis3, ZC3H18, and Ars2, and Nop60B copurified with Rrp6, Dis3, ZC3H18, and Ars2. NHP2, a component of H/ACA snoRNPs—which catalyze rRNA pseudouridylation—copurified with Rrp6, Dis3, ZC3H18, and Ars2. The nuclear exosome is required for 5.8S rRNA processing in the nucleolus(Houseley et al., 2006; Allmang et al., 2000), and M-phase phosphoprotein 6 (Mpp6),(Schilders et al., 2005) which generates 3′ ends of 5.8S rRNA, copurified with Rrp6, Dis3, and Ars2.

### Successful oogenesis requires Integrator, NEXT, and nuclear exosome components

Female flies with reduced germ-line levels of Integrator, NEXT, or nuclear exosome components were sterile (*IntS11*, *Dis3*, and *Rrp6*) or sub-fertile (*ZC3H18* and *Ars2*) (Figures 3A, 3B, and S1). We used *α-Tub67C*-Gal4 to drive germ-line expression of shRNAs targeting the mRNAs encoding the core Integrator component IntS11, the NEXT complex proteins ZC3H18 and Ars2, and the nuclear exosome proteins Dis3 and Rrp6. *α-Tub67C*-Gal4 expression begins in stage 2 egg chambers,(ElMaghraby et al., 2022) and oogenesis proceeds to at least stage 10 for all five shRNAs (Figure 4A). An shRNA targeting the eye-color gene *white* (*w*) served as control. Females depleted of *IntS11*, *ZC3H18*, *Dis3*, or *Rrp6* in the germ line had smaller ovaries than control and laid few eggs when mated to wild-type males for 10 days (Figure 3A and Figure S2). By contrast, the size and shape of ovaries in germ-line-specific *Ars2* knockdown flies were indistinguishable from wild-type. Fluorescence in situ hybridization (FISH) confirmed germ-line depletion of all five mRNAs (Figures 4, S3 and S4).

**Figure 3.**
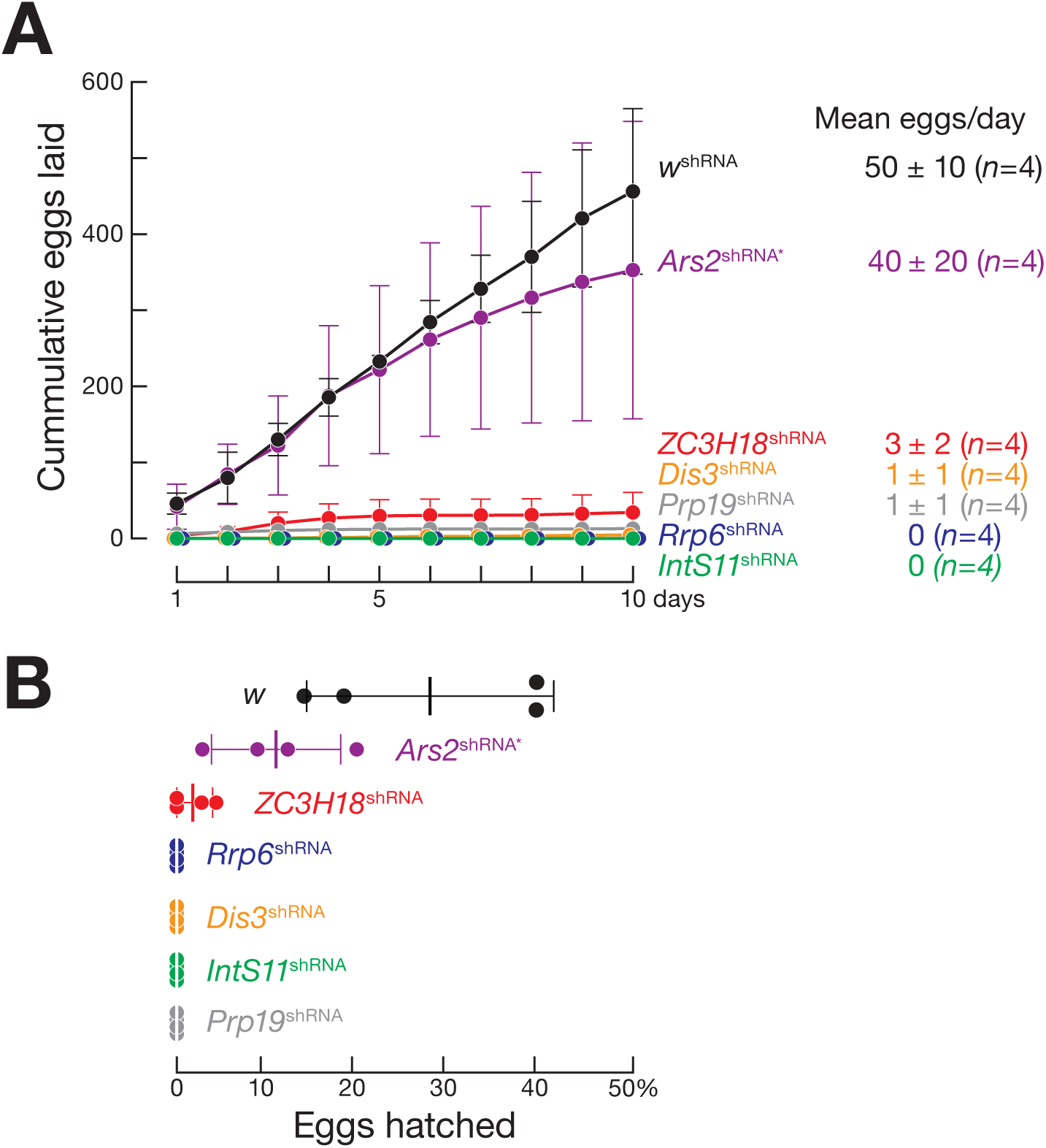
Integrator, NEXT, and nuclear exosome components are required for *Drosophila* female fertility. (A) Cumulative eggs laid in 10 days by females expressing a germ-line-specific shRNA targeting the indicated genes. Females expressing the shRNA targeting *Ars2* were heterozygous for the loss-of-function allele *Ars2^R12fsX^* (*Ars2*^shRNA*^). Females were mated to *w^1118^*males. (B) Eggs laid per day for each genotype. Vertical bars, mean; whiskers, SD (*n* = 4).

**Figure 4.**
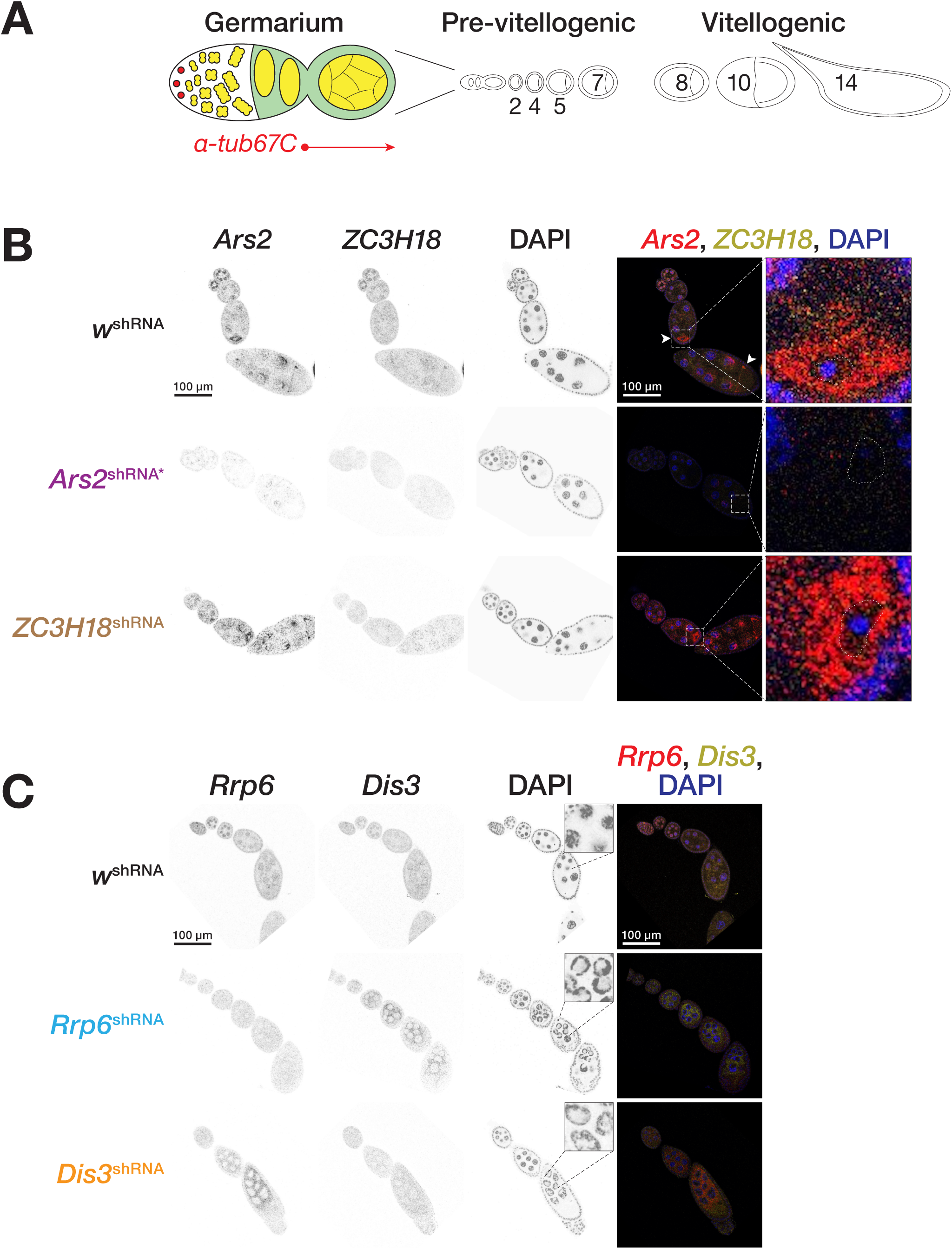
Transcriptional expression and localization of NEXT and nuclear exosome components during *Drosophila* oogenesis. (A) Schematic of *Drosophila* oogenesis. Red circles, germ-line stem cells; cysts are in yellow. (B and C) RNA fluorescent in situ hybridization (RNA FISH) of ovaries dissected from three-day-old female flies were imaged by confocal microscopy. DNA was stained with DAPI (blue). White arrowheads indicate the oocyte nucleus. Dashed white lines indicate the nuclear envelope based on wheat germ agglutinin staining (not shown). (B) RNA FISH for germ-line expression of shRNAs targeting *Ars2* or *ZC3H18*. Females expressing the shRNA targeting *Ars2* were heterozygous for the loss-of-function allele *Ars2^R12fsX^*(*Ars2*^shRNA*^). (C) RNA FISH for germ-line expression of shRNAs targeting *Rrp6* or *Dis3*. Insets show the change in nucleolar morphology when *Ars2* or *ZC3H18* was depleted.

To further decrease *Ars2*, we combined the *Ars2* shRNA with one copy of a homozygous lethal frameshift allele, *Ars2^R12fsX^* (henceforth, *Ars2*^shRNA*^). *Ars2*^shRNA*^ females laid nearly wild-type numbers of eggs: the median number of eggs per day for *Ars2 =* 40 ± 20 vs. 50 ± 10 for *w* ^shRNA^. *Ars2*^shRNA*^ ovaries were still morphologically similar to control, but fewer eggs hatched (11% ± 6 vs. 30% ± 10; *p* < 0.005, one-way ANOVA with Dunnett’s post-hoc test corrected for multiple hypothesis testing; Figure 3B).

### *Ars2* transcripts localize to the oocyte

*Ars2* transcripts localized to the nuclei of pre-vitellogenic nurse cells and later appeared in the oocyte nucleus (Figure 4A and 4B). *Ars2* was first detected in the nuclei of the germarial germ-line and follicle cells (Figure 4B and Figure S4). Between stages 5 and 7 of oogenesis, *Ars2* could be detected in both the oocyte nucleus and cytoplasm and continued to accumulate primarily in the cytoplasm until stage 9. Beginning at stage 9, Ars2 localized to the oocyte anterior (Figure 4B, white arrow). Little if any cytoplasmic *Ars2* RNA was detected in the nurse cells or oocyte in *Ars2^shRNA^*\* ovaries. However, multiple nuclear puncta remained in the nurse cell nuclei. Because RNA interference occurs in the cytoplasm, the nuclear foci likely correspond to sites of transcription. The absence of such foci in the oocyte nucleus in *Ars2^shRNA^*\* ovaries suggests that *Ars2* mRNA is produced in nurse cells and then transported to the oocyte.

### *ZC3H18* transcripts localize to both the nurse cell and oocyte nuclei

Like *Ars2*, *ZC3H18* was detected in the nuclei of germarial germ-line and follicle cells (Figure 4B and Figure S4A). Before stage 4, *ZC3H18* was concentrated in the nurse cell nuclei, but could also be detected in the cytoplasm. After stage 4, *ZC3H18* localized to the nuclei of both somatic follicle cells and nurse cells and continued to slowly increase in the cytoplasm beginning at stage 5. From stage 5 onward, *ZC3H18* was present in both the nuclei and cytoplasm of nurse cells and somatic follicle cells, persisting beyond stage 10. Germ-line depletion of *Ars2* (*Ars2*^s*hRNA*^*\**) visibly decreased the abundance of *ZC3H18* transcripts in the nucleus, suggesting that Ars2 plays a role in either the transcription or stability of nuclear *ZC3H18* RNA (Figure 4B). By contrast, the abundance and localization of *Ars2* were unaltered in the oocyte, nurse cells, or somatic follicle cells upon depletion of *ZC3H18*.

### Ars2 and ZC3H18 proteins localize to the oocyte

Ars2 and ZC3H18 proteins were both detected in the oocyte nucleus (Figure 5 and Figure S5). We used fluorescently labeled anti-FLAG antibody to detect Ars2 or ZC3H18 in homozygous epitope-tagged knock-in flies (Figure 1B). In the germarium, Ars2 localized to the nuclei of germ-line and follicle cells. During the pre-vitellogenic and vitellogenic stages of oogenesis, Ars2 was detected in the nuclei of germ-line nurse cells and somatic follicle cells, and until stage 11, the oocyte nucleus. Beginning at stage 8, Ars2 localized to the inner face of the oocyte nuclear envelope.

**Figure 5.**
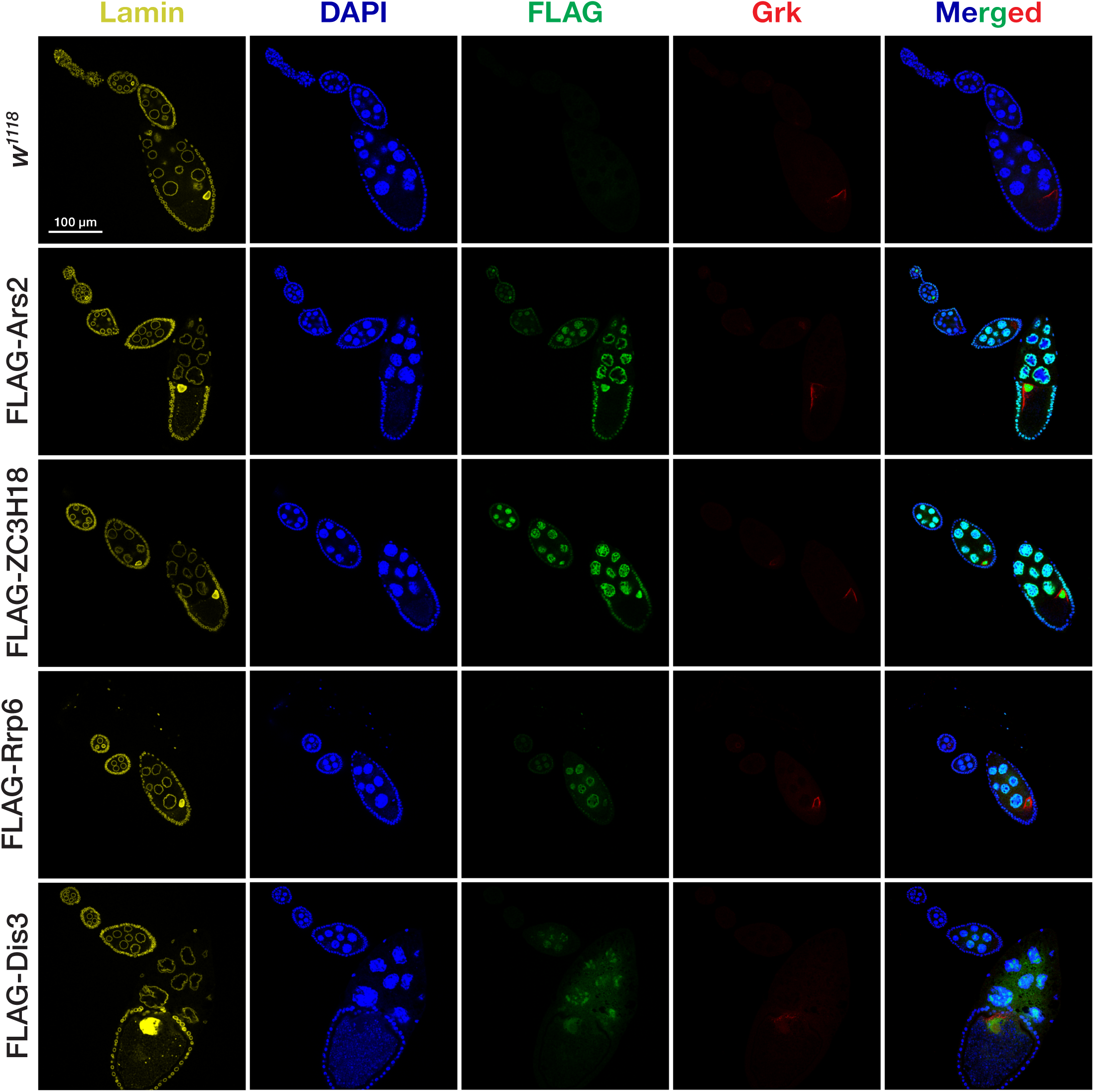
Expression and localization of epitope-tagged proteins during *Drosophila* oogenesis. Localization of 3XFLAG-epitope tagged Ars2, ZC3H18, Rrp6, and Dis3 proteins. Ovarioles are oriented to show the progression of developmental stages from top left to bottom right. Nuclei were stained with DAPI. FLAG-conjugated proteins were detected using anti-FLAG M2 antibody conjugated to CoraLite Plus 488 (green). Gurken was detected using mouse anti-Gurken antibody conjugated to CoraLite Plus 647 (red). The nuclear membrane was detected using mouse anti-Lamin antibody conjugated to CoraLite Plus 555 (yellow). *w^1118^* flies were used as a non-tagged negative control.

The localization of ZC3H18 was similar to Ars2 (Figure 5). ZC3H18 was detected in the nuclei of germarial germ-line and follicle cells (Figure S5). Starting from stage 4, ZC3H18 levels increased in both the nuclei of nurse cells and somatic follicle cells, and could be detected in stage 10 oocyte nuclei.

### *Rrp6* and *Dis3* transcripts localize to both the cytoplasm and the nucleus of nurse cells

Like *Ars2 and ZC3H18, Rrp6* and *Dis3* were detected in the nuclei of germ-line and follicle cells in the germarium (Figure 4C and Figure S4B). Furthermore, *Rrp6* and *Dis3* preferentially localized to nurse cell nuclei before stage 4, which is also similar to localization of *Ars2* and *ZC3H18* in the same stage. After stage 4, *Dis3* and *Rrp6* became readily detectable in the cytoplasm of nurse cell, while nuclear *Dis3 and Rrp6* levels slightly decreased. This pattern continued beyond stage 10.

### Depletion of *Rrp6* causes cytoplasmic accumulation of *Dis3*, and, reciprocally, *Rrp6* accumulates upon *Dis3* depletion

Germ-line depletion of *Rrp6* reduced the level of nuclear *Dis3* mRNA, yet increased the abundance of *Dis3* in the nurse cell cytoplasm (Figure 4C and Figure S4B). Conversely, germ-line depletion of *Dis3* reduced the abundance of nuclear *Rrp6* mRNA but increased *Rrp6* cytoplasmic levels.

Moreover, germ-line depletion of Dis3 or Rrp6, starting at stage 4 and persisting until oogenesis arrest at stage 9 or 10, was accompanied by altered nucleolar morphology and a visible enlargement of nurse cell nucleoli, which appeared as DAPI-negative regions within the nucleus catalytic components of exosome.

### Transcription of *Rrp6* and *Dis3* mRNAs long precedes their translation

Although *Rrp6* and *Dis3* mRNAs were readily detectable at all stages of oogenesis, Rrp6 protein was not detected before stage 5, and Dis3 was not detected before stage 7 (Figure 5 and S5). Between stages 5 and 7, Rrp6 protein localized to the nuclei and cytoplasm of both germ-line nurse cells and somatic follicle cells. In contrast, Dis3 was only detected in the nurse cell nuclei and cytoplasm, but not in follicle cells. Rrp6 or Dis3 could be weakly detected in the oocyte nucleus between stages 8 and 10, the time at which it migrates to the dorsal anterior corner of the oocyte, and *Grk* mRNA accumulates in the dorsal anterior cytoplasm adjacent to the oocyte nucleus.

### Processing of Pol II-dependent snRNAs requires Integrator and nuclear exosome components

Our data show that IntS11 establishes the 3′ ends of Pol II-dependent snRNAs during *Drosophila* oogenesis. We used Oxford Nanopore Technologies long-read sequencing of chromatin-associated, capped, non-polyadenylated ovary RNA to determine the effect of germ-line depletion of *IntS11*, *Rrp6*, *Dis3*, *ZC3H18*, or *Ars2* on co-transcriptional RNA processing. Depletion of *IntS11* or *Rrp6* caused aberrant termination or defective 3′ end processing of Pol II-dependent snRNAs but not the Pol III-transcribed 7SK snRNA (Figure 6A). Without wild-type levels of *IntS11* or *Rrp6*, transcripts from individual Pol II-dependent snRNA loci began at the annotated transcription start site but extended 0.4–3.6 kb beyond the end of the mature snRNA (Figure 6B). Germ-line depletion of *IntS11* significantly increased the length of transcripts at all nine snRNA loci examined (median increase relative to control = 1.2– 3.7, BH-corrected Mann-Whitney Wilcoxon *p* = 9.0 × 10^−54^ to 1.9 × 10^−3^; Figure S6A and Table S3). Intriguingly, germ-line depletion of the exosome component *Dis3* significantly shortened the length of snRNAs for at least six of the nine loci (median relative to control = 0.72–0.99; BH-corrected Mann-Whitney Wilcoxon *p* = 9.4 × 10^−60^ to 4.1 × 10^−2^; Figures S6A and S7 and Table S3).

**Figure 6.**
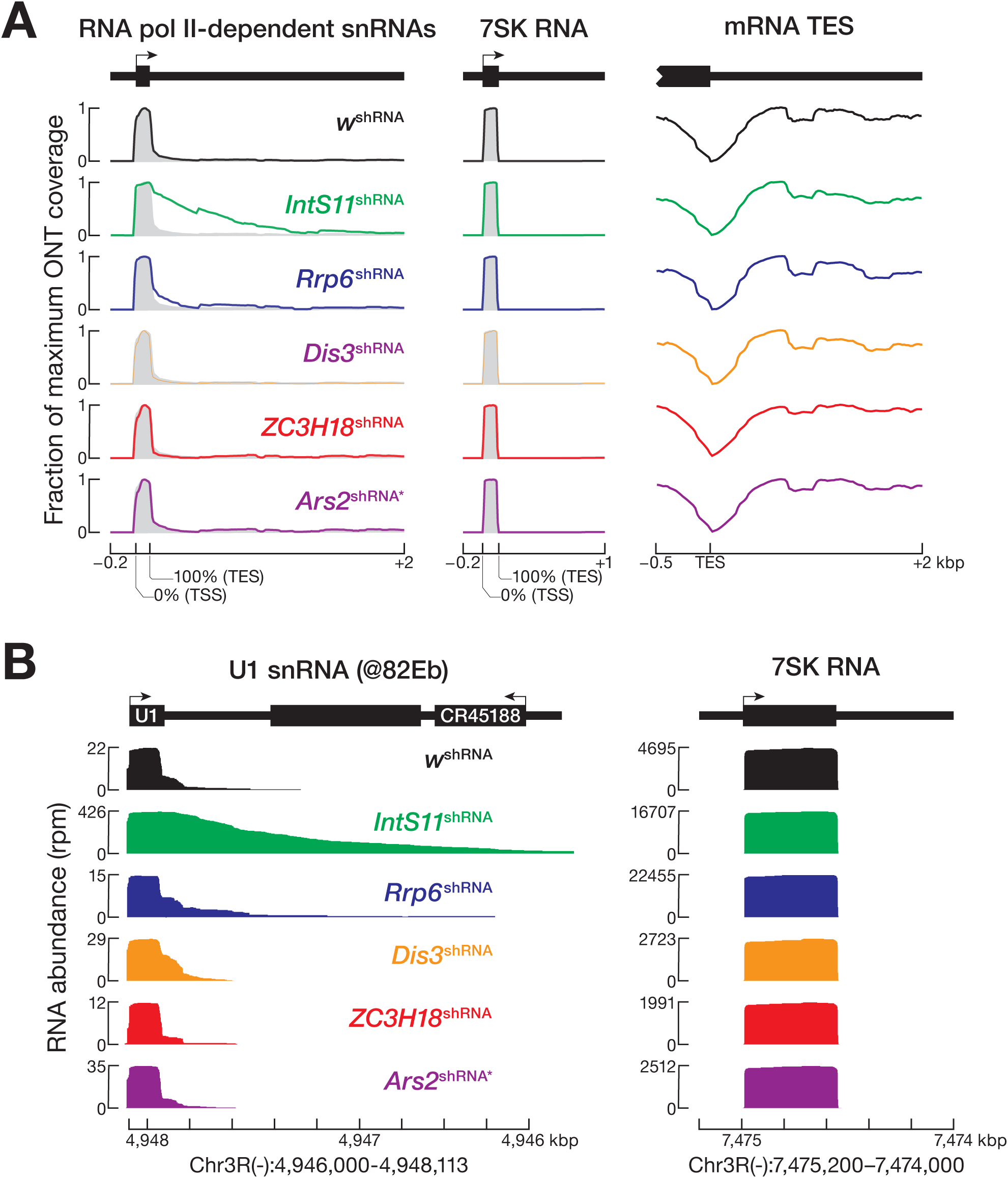
Correct 3′ end processing of Pol II-dependent snRNA transcripts requires Integrator, NEXT, and nuclear exosome. (A) Analysis of long-read sequencing of chromatin associated, non-polyadenylated ovary RNA (normalized to library sequencing depth) aligned to regions spanning 200 bp before the transcription start site (TSS) to 2000 bp after the transcription end site (TES) for 25 annotated Pol II-dependent snRNA loci, the Pol II-dependent 7SK locus, and 13,963 protein-coding genes. (B) Long-read sequencing reads (normalized to sequencing depth), aligned to the U1 snRNA locus at 82Eb and the 7SK locus.

To test whether the changes in snRNA transcript length reflected the different numbers of reads that passed our filtering criteria, we performed random sampling with replacement (bootstrapping) and compared the upper and lower 95% confidence intervals (C.I.) of each shRNA-expressing strain with the control for each snRNA locus tested (Table S4). Bootstrapping confirmed that transcript length in *IntS11*-germ-line-depleted ovaries was significantly longer and in *Dis3*-depleted germ cells was significantly shorter for all nine Pol II-dependent snRNA loci (Table S4). Finally, all of the effects of germ-line depletion of *IntS11*, *Rrp6*, and *Dis3* on snRNA biogenesis were also detected by short-read sequencing (Figures S6B, S6D and S7 and Table S5 and S6).

### Aberrant Pol II-dependent snRNA processing in fly ovaries with reduced levels of *IntS11* perturbs pre-mRNA splicing

We used short-read RNA-seq data to measure the effect of germ-line depletion of *IntS11* on pre-mRNA splicing. For each splice site, we measured the abundance of RNA-seq reads containing an exon-intron or intron-exon junction compared to the total of all splice site reads (i.e., exon-exon, exon-intron, and intron-exon junctions). Compared to control ovaries, germ-line expression of an shRNA targeting *IntS11* increased the median fraction of splice sites that failed to be spliced 2.4-fold (Benjamini-Hochberg [BH] corrected Dunn’s post-hoc test *p* = 8.8 × 10^−73^) (Figure 7A). For comparison, the germ-line depletion of the essential splicing factor Prp19 increased the median abundance of unspliced reads by 1.7-fold (BH-corrected Dunn’s post-hoc test *p* = 1.2 × 10^−34^).(Chan et al., 2003; Tarn et al., 1994) Germ-line depletion of *Prp19* yielded ovaries of normal size, but females laid fewer eggs compared to the control and none of the eggs hatched (Figure 3A and 3B).

**Figure 7.**
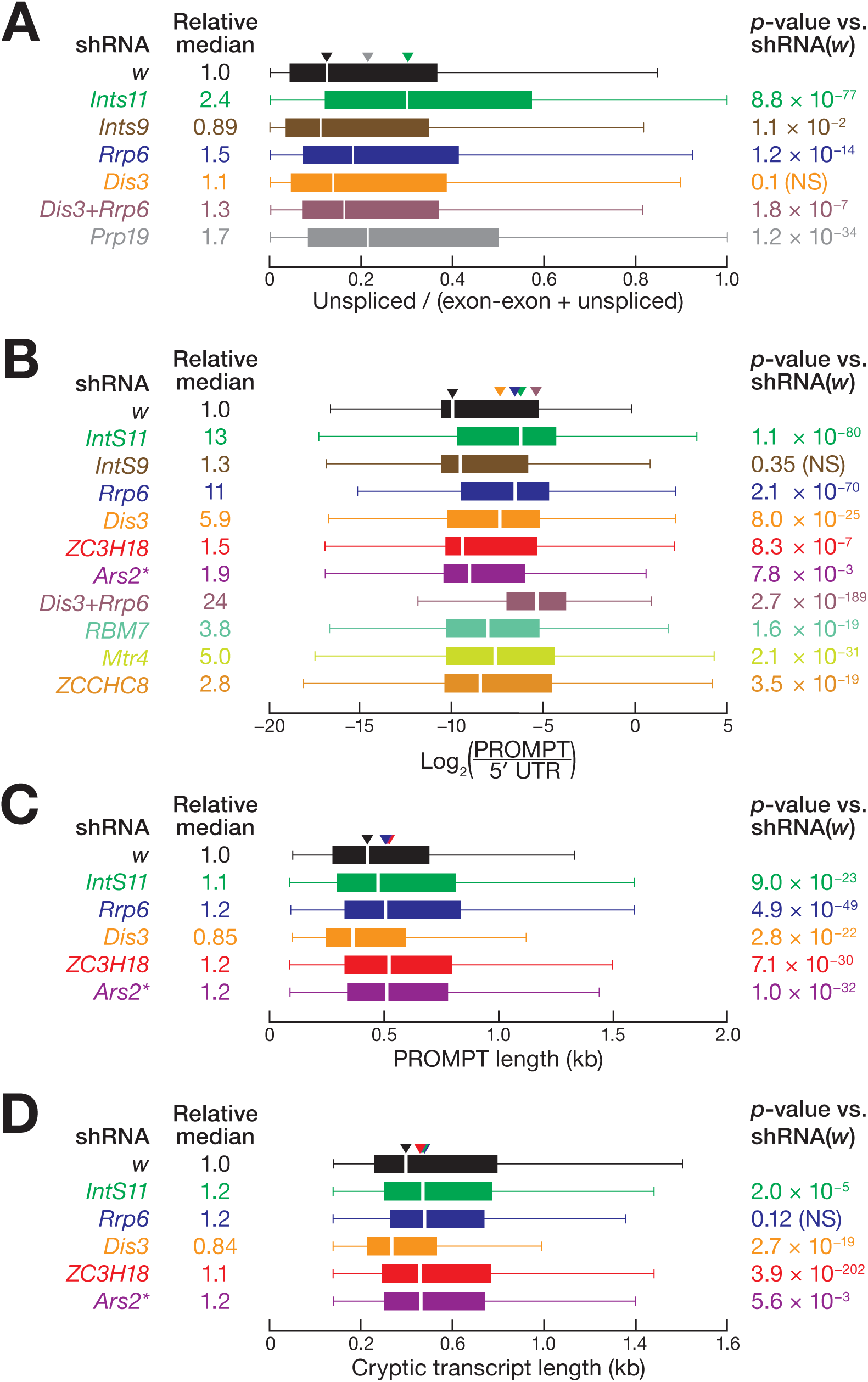
Integrator and NEXT exosome-related factor regulate thousands of cryptic loci in the *D. melanogaster* ovary. (A) Fraction of unspliced reads for the sum of exon-intron, intron-exon, and exon-exon reads for 2,509 splice sites. (B) Short-read RNA sequencing abundance for 2,527 distinct PROMPT regions relative to the divergently transcribed 5′ UTR. *RBM7*, *Mtr4*, and *ZCCHC8* were depleted using long-hairpin RNA instead of an shRNA. (C) Lengths of long-read sequences whose 5′ ends were in annotated PROMPT regions. Total number of reads analyzed for each genotype: *w*^shRNA^ = 4,429; *IntS11*^shRNA^ *=* 18,678; *Rrp6*^shRNA^ = 11,733; *Dis3*^shRNA^ = 4,887; *ZC3H18*^shRNA^ = 4,205, and *Ars2*^shRNA*^ = 5,688. (D) Lengths of long-read sequences from 1,140 unannotated genomic regions predicted by GRO-seq data to be transcriptionally active. Boxplots depict the inter-quartile ranges, median, maximum and minimum values after excluding outliers; BH-corrected Dunn’s post-hoc test *p*-values are shown.

### The Integrator, NEXT, and nuclear exosome complexes prevent the accumulation of PROMPT and cryptic transcripts in vivo

Germ-line depletion of Integrator, NEXT and nuclear exosome complex components in the *Drosophila* female germ-line recapitulates the accumulation of aberrant transcripts upon depletion of these complexes in *Drosophila* and human cell lines.(Rennie et al., 2018; Szczepińska et al., 2015; Lubas et al., 2011) Divergent transcription of the DNA strand opposite annotated genes naturally produces promoter upstream transcripts (PROMPTs), which are then rapidly degraded by Integrator or directed by the NEXT complex to the nuclear exosome.(Yang et al., 2024; Rennie et al., 2018) We used short-read RNA sequencing of whole ovaries to measure the steady-state abundance of PROMPTs in flies depleted of Integrator, NEXT, or nuclear exosome components in the female germ line. To identify potential PROMPT-producing loci, we examined the 2,000 bp upstream of the transcription-start sites of protein coding genes to detect divergent transcripts templated by the opposite DNA strand. In order to exclude loci for which the mRNA-producing promoter was not responsible for the production of the putative PROMPT, we considered only loci for which the expression of the divergent transcript was less than that of the corresponding mRNA in the control ovaries. To account for differences in promoter activity among the loci, we calculated the ratio of the abundance of each PROMPT, divided by 2000, to the length-normalized abundance of the 5′ UTR of divergently transcribed mRNA. Analysis of the resulting 2,527 loci revealed significantly more PROMPT production in ovaries depleted of the Integrator complex component *IntS11*, the NEXT complex components *ZC3H18*, *Rbm7*, *Mtr4,* or *ZCCHC8*, and the nuclear exosome components *Rrp6* and *Dis3* (median increase relative to control = 1.3–13; BH-corrected Dunn’s post-hoc test *p* = 2.7 × 10^−189^–7.8 × 10^−3^; Figure 7B). Notably, the abundance of PROMPTs was highest when *Dis3* and *Rrp6* were simultaneously depleted using germ-line-expressed shRNAs, consistent with the idea that the combined action of the two nuclear exosome ribonucleases normally prevents the accumulation of these RNAs.

We used long-read sequencing of chromatin-associated RNA to compare the length of PROMPTs—regardless of abundance—in the shRNA-expressing ovaries to the control. PROMPT length was significantly longer when *IntS11*, *Ars2*, *ZC3H18*, or *Rrp6* were depleted in the germ line (median increase relative to control = 1.1–1.2; BH-corrected Dunn’s post-hoc test *p* = 1.0 × 10^−32^–9.0 × 10^−23^; Figure 7C). By contrast, PROMPTs were significantly shorter in *Dis3*-depleted germ cells (median decrease relative to control = 0.85, BH-corrected Dunn’s post-hoc test *p* = 3.4 × 10^−24^; Figure 7C). Long-read sequencing also revealed cryptic transcripts from unannotated but transcriptionally active regions of the genome. These cryptic transcripts were significantly longer in the ovaries of flies expressing in the germ-line shRNAs targeting *IntS11*, *Ars2*, *ZC3H18*, or *Rrp6* (median increase relative to control = 1.1–1.2, BH-corrected Dunn’s post-hoc test *p* = 3.9 × 10^−202^–2.0 × 10^−5^), but shorter in *Dis3*-depleted ovaries (median decrease relative to control = 0.84; BH-corrected Dunn’s post-hoc test *p* = 1.4 × 10^−56^; Figure 7D). Although PROMPTs and intergenic cryptic transcripts are generated from different genomic locations, both types of aberrant RNA accumulated in the absence of wild-type levels of IntS11, Ars2, ZC3H18, or Rrp6. Our data suggest that suppression of both PROMPTs and cryptic transcripts requires the combined action of the Integrator, NEXT, and nuclear exosome complexes.

## DISCUSSION

### DTME crosslinking with SDS improves protein complex purification

To identify proteins interacting with the Integrator, NEXT, and nuclear exosome complexes in *Drosophila*, we crosslinked ovaries with DTME, a cleavable sulfhydryl crosslinker, and identified the key proteins copurifying with each complex. In preliminary experiments using non-denaturing buffers, the yield of copurifying proteins was low and the background high. The addition of 0.5% SDS substantially improved the yield of both the epitope-tagged protein and its copurifying interactors. Moreover, in the presence of 0.5% SDS most proteins that copurified with the tagged bait were undetectable in the control purification using ovaries from non-tagged flies: 73% of proteins copurifying with tagged IntS11 were undetectable in the control (581/797); 71% for Rrp6 (537/753); 76% for Dis3 (676/892); 75% for ZC3H18 (633/849); and 76% for Ars2 (698/914).

### Crosslinking and purification of the IntS11 catalytic nuclease in the *Drosophila* germ line recapitulates previously identified interactors

The Integrator complex comprises 15 core proteins.(Fianu et al., 2024; Zheng et al., 2023; Azuma et al., 2023; Offley et al., 2023; Baillat et al., 2005; Chen et al., 2012) Five of these proteins were identified as interacting with IntS11 in *Drosophila* ovaries using a novel in vivo crosslinking and copurification strategy: IntS1, IntS4, IntS9, Ints12, and IntS13. IntS9 has been shown to form a stable dimer with IntS11; IntS4 binds this dimer in the nucleus to form the Integrator cleavage module.(Albrecht and Wagner, 2012; Albrecht et al., 2018; Pfleiderer and Galej, 2021; Wu et al., 2017) The protein BRAT1 inhibits the IntS11 endonuclease in the cytoplasm and escorts the IntS9•IntS11 dimer to the nucleus(Cihlarova et al., 2022; Dokaneheifard et al., 2024; Sabath et al., 2024) BRAT1 was the most abundant IntS11 interactor in our experiments.

Two lines of evidence argue that a substantial fraction of the Integrator complex we isolated is either free or actively terminating transcription: (1) we failed to recover DSS1, IntS3, IntS5, and IntS7, components associated with the post-termination Integrator complex(Fianu et al., 2024); and (2) the PP2A subunits Tws and Wrd(Janssens and Goris, 2001), which are not present in the post-termination complex(Fianu et al., 2024), both copurified with IntS11. During *Drosophila* oogenesis, the nurse cell DNA undergoes endoreduplication (16C) to support the massive transcriptional load required to supply the egg with maternal RNA and protein to support the early phase of embryo development. Splicing all the pre-mRNA produced in the nurse cells likely requires high levels of snRNAs, who maturation would in turn require both high levels and rapid recycling of Integrator.

Our findings also suggest that IntS11 may collaborate with the 19S proteasome complex. In the nucleus, the 19S proteasome regulatory particle has been implicated in both proteolytic and non-proteolytic roles in transcription.(Arkinson et al., 2025; Tomko and Hochstrasser, 2011; von Mikecz, 2006) IntS11 specifically copurified with eight subunits of the 19S proteasome control: Rpn1–3, Rpn5, Rpn8, and Rpn10–12. Modeling with AlphaFold multimer predicts that Rpn2, Rpn3, Rpn8, and Rpn10 bind directly to IntS11 (MCS >0.5). We speculate that the Integrator complex collaborates with the nuclear proteasome through direct protein-protein interactions, perhaps to coordinate the destruction of RNA and protein in aberrant RNPs or regulate transcription rates. Alternatively, the association of IntS11 with the proteasome may simply reflect degradation of improperly assembled or free IntS11 to prevent it from non-specifically degrading RNAs.

### Depletion of Integrator impairs snRNA biogenesis and leads to the accumulation of aberrant transcripts

Depletion of Integrator subunits causes accumulation of unprocessed snRNAs in human cell lines, *Caenorhabditis elegans*, and planaria.(Gómez-Orte et al., 2019; Schmidt et al., 2018; O’Reilly et al., 2014) Similarly, depletion of *IntS11* in the female fly germ line led to accumulation of improperly processed snRNA transcripts with long 3′ extensions. The defect in snRNA processing was accompanied by an increase in unspliced mRNAs in *IntS11-*depleted fly ovaries. A second function of the Integrator complex is to deliver PP2A to elongating Pol II to terminate production of aberrant transcripts.(Vos et al., 2018; Sun et al., 2010; Cortazar et al., 2019; Egloff et al., 2010) Similarly, our data from *Drosophila* ovaries show that depletion of *IntS11* leads to accumulation of PROMPTs as well as cryptic transcripts from unannotated regions of the genome, underlining the critical contribution of the Integrator complex in nuclear RNA homeostasis during *Drosophila* oogenesis.

### Ars2 and ZC3H18 connect NEXT to nuclear exosome *Drosophila* ovaries

In the fly ovary, epitope-tagged Ars2 and ZC3H18 copurified with each other, with other components of the NEXT complex, and with components of the nuclear exosome complex. In mammalian cultured cells, binding of ARS2 and ZC3H18 to the cap-binding complex directs nascent transcripts to the nuclear exosome through binding of the NEXT complex component MTR4 with the exosome nuclease RRP6.(Andersen et al., 2013; Gerlach et al., 2022; Puno and Lima, 2018; Puno and Lima, 2022) Similarly, both Ars2 and ZC3H18 copurified with Nuclear Cap-Binding Protein 1 (NCBP1) and Mtr4. Interestingly, neither epitope-tagged Ars2 nor ZC3H18 copurified with the *Drosophila* homolog of RBM7; in mammals RBM7 binds transcripts with 3′ poly(U) stretches, enabling NEXT to target them for destruction by the nuclear exosome.(Falk et al., 2016) Whether RBM7 plays a similar role in flies remains to be determined.

### Ars2 and ZC3H18 bind the PP2A complex in the fly ovary

The *Drosophila* PP2A complex comprises a structural A subunit, PP2A-A, three regulatory B subunits, Twins (Tws), Widerborst (Wdb), and Well rounded (Wrd), and a catalytic C subunit, Microtubule star (Mts).(Janssens and Goris, 2001) Our combined LC/MS data and modeling analyses suggest that PP2A-A and Tws can directly bind Ars2 and ZC3H18. To our knowledge, an association of PP2A with the NEXT complex has not been previously reported. The Integrator complex is unlikely to mediate the interaction between PP2A and Ars2 or ZC3H18: only a single Integrator complex component (IntS13) copurified with Ars2 and none copurified with ZC3H18. Structural analyses of Integrator with PP2A show that while IntS1, IntS2, and IntS5–IntS8 bind directly to surfaces on PP2A, IntS13 does not.(Fianu et al., 2024; Zheng et al., 2020) In human cells, INTS13 may contribute to monocytic/macrophagic differentiation through a pathway that does not rely on the Integrator catalytic subunit INTS11: we speculate that the copurification of fly IntS13 with Ars2 and ZC3H18 reflects a NEXT complex pathway involving IntS13 but not other core Integrator complex subunits.(Barbieri et al., 2018)

### A potential function of the NEXT and nuclear exosome complexes during transcriptional reactivation of the *Drosophila* oocyte

Unlike the nurse cells, the oocyte nucleus is transcriptionally quiescence until stage 9, when chromatin remodeling reactivates transcription of its diploid genome.(Navarro-Costa et al., 2016) Yet only a small number of RNAs have been demonstrated to localize to the oocyte(Jambor et al., 2015) when transcription is reactivated, and only one—*Grk*—has been shown to be transcribed from the oocyte genome.(Saunders and Cohen, 1999) Our finding that the NEXT complex components Ars2 and ZC3H18 and the exosome nucleases Dis3 and Rrp6 accumulate in the oocyte nucleus at stage 9 suggests that they may collaborate to degrade RNA transcripts produced when the oocyte genome undergoes chromatin remodeling.

### The two catalytic subunits of the nuclear exosome, Rrp6 and Dis3, collaborate to degrade aberrant transcripts

The RNA exosome exists in cytoplasmic and nuclear forms.(Allmang et al., 1999; Tomecki et al., 2010; Kilchert et al., 2016) Rrp6 is restricted to the nuclear exosome, whereas Dis3 can be detected in both the cytoplasmic and nuclear complexes in *Drosophila* cell lines.(Mitchell et al., 1997; Graham et al., 2006) We detected Rrp6 predominantly in nurse cell nuclei, whereas Dis3 was present in both the nuclei and cytoplasm. Depletion of *Rrp6* from fly germ-line cells caused a larger accumulation of PROMPTs than depletion of *Dis3.* When both *Rrp6* and *Dis3* were depleted, PROMPTs abundance more than doubled, compared to depletion of Rrp6. These findings suggest that the combined action of Rrp6 and Dis3 is required to fully suppress accumulation of PROMPT RNAs at bidirectionally transcribed genes in the female fly germ line.

### A resource for investigating nuclear homeostasis in vivo

Here we describe a novel in vivo crosslinking/copurification/mass spectrometry workflow that allowed us to determine the interactome of key components of the Integrator, NEXT, and nuclear exosome complexes in the *Drosophila* ovary. Combining this experimental method with modeling using AlphaFold allowed us to rank the likelihood that individual proteins bind directly to the epitope-tagged bait. Once a protein of interest is epitope tagged by standard knock-in methods, our method can be used to define the high-confidence interactome of any cellular complex. To assess the in vivo impact on oogenesis of depletion of individual Integrator, NEXT, and nuclear exosome components, we developed a novel long-read ONT-based method to sequence chromatin-associated, capped, non-polyadenylated RNA. We used this method to survey the landscape of RNAs normally processed by Integrator, NEXT, and nuclear exosome complexes. Together with high-depth short-read sequencing, our data sets provide a resource for dissecting the in vivo role of the Integrator, NEXT, and nuclear exosome complexes in maintaining nuclear RNA homeostasis during *Drosophila* oogenesis.

### Limitations of this study

Crosslinking using DTME in our mass spectrometry analysis covalently links sulfhydryl groups between proteins in the range of 13.3 Å. If proteins do not contain appropriately placed cysteines, they will fail to crosslink to the tagged protein. Using crosslinking agents reacting with other chemical moieties could allow detection of additional proteins.

## RESOURCE AVAILABILITY

### Lead contact

Further information and requests for resources and reagents should be directed to, and will be fulfilled by, the Lead Contact, Phillip D. Zamore or by completing the request form at https://www.zamorelab.umassmed.edu/reagents.

### Materials availability

Reagents generated in this study are available from the Lead Contact.

### Data and code availability

Proteomics data are available from Mass spectrometry Interactive Virtual Environment MSV000096612 (ftp://MSV000096612@massive.ucsd.edu/). Sequencing data are available from the National Center for Biotechnology Information Sequence Read Archive using accession number PRJNA1185634. All software used for the analyses in this paper are detailed in the STAR Methods. All information required for data analysis in this paper will be made available upon request to the lead contact.

## ACKNOWLEDGMENTS

We thank Beatrix Ueberheide and members of the New York University Proteomics Laboratory for assistance with Mass spectrometry, Birgit Koppetsch for imaging assistance, and members of Zamore laboratory for help, advice, discussions, and comments on the manuscript. This work was supported in part by NIH grant R35 GM136275 to P.D.Z.

## AUTHOR CONTRIBUTIONS

Conceptualization, Y.L. and P.D.Z.; methodology, Y.L.; experiments, Y.L.; fly husbandry and dissection, Y.L. and C.T.; imaging, Y.L.; Imaging analysis, Y.L.; LC-MS/MS data analysis, Y.L. and A.B.; RNA sequencing data processing and analysis, A.B. and S.H.H.; writing – original draft, Y.L. and A.B.; writing – review & editing, P.D.Z., Y.L., and A.B.; supervision and funding acquisition, P.D.Z.

## STAR METHODS

### Key Resources Table

#### EXPERIMENTAL MODEL

Flies were grown at 25°C. shRNA lines targeting *w* (stock #33613), *IntS11* (#65093), *Rrp6* (#34809), *Dis3* (#67919), *ZC3H18* (#55353), *Ars2* (#35204), *Prp19* (#32865), and *IntS9* (#65892) were from the Bloomington Drosophila Stock Center. α-Tubulin67C-GAL4 (#7063) was used to drive short hairpin RNA transcription in the female germ line. Long-hairpin RNA-expressing lines targeting *Mtr4* (#108847), *ZCCHC8* (#106312), and *RBM7* (#451118) were from the Vienna Drosophila Resource Center (VDRC) and co-expressed with UAS-*dicer-2* under the control of α-Tubulin67C-GAL4 driver.

### Generation of *Ars2* frameshift mutant

Two guide RNA sequences targeting the first exon of *Ars2* were cloned into pCFD4d, and the plasmid was injected into *vas*-Cas9 fly embryos (Bloomington Drosophila Stock Center, 51323, Rainbow Transgenic Flies, Camarillo, CA, USA).(Port et al., 2014). From these flies, a line, *Ars2^R12fsX^*, was isolated that contained a single-nucleotide deletion at chr2R:6,085,192 (*D. melanogaster* genome release r6.07) that causes a heterozygous viable, homozygous lethal frameshift introducing 102 new amino acids followed by a premature stop codon. The *Ars2^R12fsX^*mutant was balanced against *CyO* and backcrossed to *w^1118^*for 10 generations.

### Epitope-tagged flies

sgRNA and donor DNA were cloned into plasmid pCFD3 and pBluescript II KS, respectively, and injected into *vas*-Cas9 fly embryos (Bloomington Drosophila Stock Center, 51324, CRISPR Core, University of Massachusetts Chan Medical School). Donor sequences introduced Twin-Strep and 3XFLAG epitope tags immediately upstream of the start codon of each gene and contained >500 bp of homologous sequence for each arm. For *IntS11,* the donor encompassed 1306 bp spanning genomic nucleotides chr3R:29,744,985–29,746,036; for *Rrp6*, 1337 bp spanning genomic nucleotides chr3R:14,885,417–14,886,497; for *Dis3*, 1351 bp spanning genomic nucleotides chr3R:24,332,931–24,334,025; for *ZC3H18*, 1332 bp spanning chrX:7,284,204–7,285,272; for *Ars2*, 1316 bp spanning genomic nucleotides chr2R:6,084,691–6,085,750. For each gene, two sgRNAs were cloned into pCFD3.(Port et al., 2014) Donor and guide plasmids were co-injected into *vas*-Cas9 embryos. Knock-in flies were backcrossed to *w*^1118^ for at least six generations. Table S7 lists sgRNA, donor DNA, and genotyping primer sequences.

### Genotyping

Flies were ground with a plastic pestle in squishing buffer (10 mM Tris-Cl, pH 8.0, 1 mM ethylenediaminetetraacetic acid (EDTA), 25 mM NaCl, 200 µg/ml freshly diluted Proteinase K; Thermo Fisher Scientific).(Ge et al., 2016) Homogenized lysate was incubated at 55°C for one hour. Lysate was centrifuged to precipitate the cell debris, and the supernatant was used as PCR template. PCR amplification was performed using MeanGreen 2×Taq master mix (Empirical Bioscience) with 10 µM each primer under the following conditions: 95°C for 30 sec, 35 cycles of 95°C for 30 sec, 65°C for 30 sec, 72°C for 60 sec, final extension at 72°C for 5 min. Primer sequences are listed in Table S7. Electrophoresis was performed in a 2% (w/v) agarose gel stained with 0.5 μg/ml ethidium bromide.

### Crosslinking and Twin-Strep tag purification

Two hundred female flies expressing amino-terminally tagged IntS11, Rrp6, Dis3, ZC3H18, or Ars2 were collected 2 days after eclosion and fed yeast for 3 days. *w^1118^* flies were used as a non-tagged control. Ovaries were dissected in phosphate buffered saline (PBS; 137 mM sodium chloride, 10 mM disodium phosphate, 2.7 mM potassium chloride, 1.8 mM monopotassium phosphate, pH 7.4), washed twice with cold PBS, and chemically crosslinked with 5 mM dithio-bis-maleimidoethane (DTME) (Thermo Fisher Scientific) in 500 μl PBS per two hundred pairs of ovaries at 25°C for 15 min. Crosslinked ovaries were then washed three times with PBS for 10 min each and homogenized by a Dounce homogenizer with 30 strokes of a type B pestle in lysis buffer containing 0.5% (w/v) sodium dodecyl sulfate (SDS), 50 mM HEPES-KOH, (pH 7.6), 100 mM potassium acetate, 50 mM L-Arginine, 2.5 mM magnesium acetate, 0.1% (v/v) Tween-20, 1 mM DTT, 10 U Turbo DNase (Thermo Fisher Scientific), 25 U RNase A/T1 (Thermo Fisher Scientific), and protease inhibitor cocktail (1 mM AEBSF (4-(2-aminoethyl) benzenesulfonyl fluoride hydrochloride [EMD Millipore], 0.3 μM Aprotinin [Bio Basic], 20 μM Bestatin [Sigma Aldrich], 10 μM E-64 [(1S,2S)-2-(((S)-1-((4-Guanidinobutyl)amino)-4-methyl-1-oxopentan-2-yl)carbamoyl)cyclopropanecarboxylic acid; VWR #97063], 10 μM Leupeptin [Fisher Scientific #108975]). While cooled in ice-water, the lysate was sonicated (Sonic Dismembrator Model 120, Fisher Scientific) six times for 20 sec at 35% amplitude every 2 min. Ovary lysate was centrifuged at 20,000 × *g* for 20 min at 4°C and the supernatant collected. To purify Twin-Strep-tagged proteins, MagStrep Strep-Tactin XT beads (IBA Lifesciences) were washed with lysis buffer three times, and then incubated for 30 min with lysate at room temperature, rotating. Beads were collected in a magnetic stand, then washed twice with Tris-buffered saline (TBS, 150 mM sodium chloride, 50 mM Tris, pH 7.6) containing 1 mM EDTA. Proteins were eluted by incubating the beads for 1 h with 200 mM Tris-Cl, 300 mM NaCl, 2 mM EDTA, 100 mM biotin) at 4°C, rotating.

### Mass spectrometry

#### Protein digestion

Sample preparation and analysis for mass spectrometry were performed at the New York University Langone Proteomics Laboratory. Affinity-tag purified proteins were reduced with 5 mM DTT at 57°C for 1 h, alkylated with iodoacetamide (IAA) at room temperature for 1 h, and digested with trypsin (sequencing-grade, Promega) overnight at 37°C. Digested peptides were acidified with 0.2% (v/v) trifluoroacetic acid and desalted on a C18 spin column. The eluate was lyophilized (SpeedVac) and then resolubilized in 2% acetonitrile, 0.5% acetic acid and analyzed by LC-MS/MS.

#### LC-MS/MS

LC separation was performed online on EASY-nLC 1200 (Thermo Scientific) using an Acclaim PepMap 100 pre-column (75 µM × 2 cm) and PepMap RSLC C18 analytical column (2 µM, 100 Å, 75 µM × 50 cm). Peptides were gradient-eluted from the column directly into an LTQ Orbitrap Eclipse mass spectrometer (Thermo Fisher Scientific) with Solvent A (2% acetonitrile, 0.5% acetic acid) and Solvent B (80% acetonitrile, 0.5% acetic acid) at 200 nl/min flow rate. High-resolution full MS spectra were acquired with a resolution of 120,000, an AGC target of 4 × 10^5^, with a maximum ion injection time of 50 ms and a scan range of 400–1500 m/z. Following each full MS scan, twenty data-dependent high resolution HCD MS/MS spectra were acquired. MS/MS spectra were collected using the following instrument parameters: resolution of 30,000, AGC target of 2 × 10^5^, maximum ion time of 200 ms, one microscan, 2 m/z isolation window, a normalized collision energy (NCE) of 27, and dynamic exclusion for 30 sec. Both MS and MS spectra were recorded in profile mode.

#### LC-MS/MS data analysis

MS data were analyzed using MaxQuant software version 1.6.3.4 and searched against the SwissProt subset of the Drosophila melanogaster UniProt database (http://www.uniprot.org/). (Cox and Mann, 2008) Database search was performed in Andromeda integrated in MaxQuant environment.(Cox et al., 2011) A list of 248 common laboratory contaminants included in MaxQuant was also added to the database as well as reversed versions of all sequences. For searching, the enzyme specificity was set to trypsin with the maximum number of missed cleavages set to two. The precursor mass tolerance was set to 20 ppm for the first search used for non-linear mass re-calibration and then to 6 ppm for the main search.(Cox et al., 2011) Oxidation of methionine was searched as a variable modification; carbamidomethylation of cysteines was searched as a fixed modification. The false discovery rate (FDR) for peptide, protein, and site identification was set to 1%, the minimum peptide length was set to six. Normalized label-free quantification (LFQ) intensity values (Log_2_) were used to measure protein abundance. Unified PolySTest, which integrates LIMMA, Miss test, rank products, and permutation tests, was used to obtain false discovery rates.(Schwämmle et al., 2020) LFQ intensity values were compared to the non-tagged control, and proteins enriched in the control were excluded.

#### AlphaFold Multimer

Proteins copurifying with IntS11 (460), Rrp6 (107), Dis3 (109), ZC3H18 (109), and Ars2 (213) were evaluated for their predicted ability to bind each target protein. To evaluate the accuracy of its predictions, AlphaFold Multimer provides the modeling confidence score (MCS), the weighted sum of the predicted template modeling (pTM) score and the interface predicted template modeling (ipTM): (0.8 × ipTM) + (0.2 × pTM).(Evans et al., 2021) AlphaFold2.3.1 was used for prediction, with the maximum template release date fixed as of June 18, 2024. Protein sequences were obtained from UniProt.

#### Female fertility

Female fertility was assayed as described with modifications.(Li et al., 2009) VALIUM (Vermilion-AttB-loxP-Intron-UAS-MCS) flies expressing gene-specific shRNAs were crossed to α*-tub67C GAL4* driver at 25°C. Female progeny (F1) flies were collected <1 day after eclosion. A single F1 female and three *w^1118^*male flies were mated and grown on 60 mM diameter grape juice agar plates with a blob of yeast paste (Lesaffre Red Star bakers active dry yeast) at 25°C, covered with small cages. Plates were removed and replaced every 24 h. The number of eggs laid was counted, the plate was further incubated at 25°C for 24 h, and then the number of hatched eggs was counted. Plates were scored daily for ten consecutive days. Fertility assays were performed in quadruplicate.

#### RNA fluorescence in situ hybridization (FISH)

A set of Stellaris singly-labelled 48 probes (LGC Biosearch Technologies) was used to detect transcripts (Table S8). One-day-old female flies were fed yeast paste for 2 days before 10–15 pairs of ovaries were dissected, washed twice with PBS, then fixed in PBS containing 500 μl 4% (v/v) methanol-free formaldehyde and 0.15% (v/v) Triton X-100 at room temperature for 30 min with gentle agitation. Fixed ovaries were washed three times with 0.3% (v/v) Triton X-100 in PBS and permeabilized with 500 μl 70% ethanol at 4°C overnight with rotation. After permeabilization, ovaries were rehydrated in 500 μl Stellaris RNA FISH Wash Buffer A (LGC Biosearch Technologies) containing 10% deionized formamide at room temperature for 5 min. Stellaris probes (250 M f.c.) in 50 μl Stellaris RNA FISH hybridization buffer (LGC Biosearch Technologies) containing 10% deionized formamide were added to the fixed ovaries and incubated at 37°C, rotating, overnight in the dark. Fixed ovaries were briefly rinsed with Wash Buffer A, then washed twice at 37°C for 15 min. To detect membranes, ovaries were incubated with 5 ng/µl Wheat Germ Agglutinin Alexa 488 (Thermo Fisher Scientific) at room temperature for 1 h, then washed with 500 µl of PBS containing 0.3% Triton X-100 (PBX) three times. Ovaries were incubated with 0.5 μg/ml 4′,6-diamidino-2-phenylindole (DAPI) at room temperature for 15 min, rotating in the dark. Ovaries were then washed with Wash Buffer B (LGC Biosearch Technologies) at room temperature for 5 min. Ovaries were mounted on glass slides in VECTASHIELD mounting medium (Vector Laboratories) using 22 × 40 mM cover slips and sealed with nail polish. RNA FISH images were acquired on a SP-8 Leica confocal microscope using a 40 × HC PL APO CS2 oil immersion objective (NA 1.3) and 1–2 µM-thick z-stacks. For *Ars2* and *ZC3H18* transcripts, Z-stacks were summed as follows: *w^shRNA^*, 25 1.5-µm stacks; *Ars2^shRNA^*\*, 20 2-µm stacks; *ZC3H18^shRNA^*, 20 2-µm stacks. For *Rrp6* and *Dis3* transcripts, Z-stacks were summed as follows: *w^shRNA^*, 25 1.6-µm stacks; *Rrp6^shRNA^*, 20 1.7-µm stacks; *Dis3^shRNA^*, 20 1.2-µm stacks. For *IntS11* transcript, 25 1.7-µm z-stacks were summed for *w*^shRNA^; 20, 1.3-µm z-stacks were summed for *IntS11*^shRNA^.

#### Immunofluorescence

Ovaries from 3–5-day-old flies fed yeast for 2 days were dissected, teased apart into individual ovarioles, then washed three times with PBS. Ovaries were fixed in 500 μl 4% (v/v) methanol-free formaldehyde at room temperature for 15 min with gentle agitation, then washed with PBS containing 0.1% Triton X-100 (PBX). Samples were pre-incubated with mouse serum (Sigma Aldrich M5905) at room temperature for 1 h. Antibodies were directly labeled with fluorophores (CoraLite Plus Fluorescent dyes, Proteintech) according to the manufacturer’s instructions. Mouse anti-FLAG antibodies (M2, Sigma) were conjugated to CoraLite Plus 488 dye and used at 1:1000 dilution. Mouse anti-Lamin antibodies (ADL84.12, Developmental Studies Hybridoma Bank) were conjugated to CoraLite Plus 555 and used at 1:400. Mouse anti-Gurken was conjugated to CoraLite Plus 647 and used at 1:100. Ovary samples were incubated in the dark with dye-labeled antibodies overnight at 4°C with rotation, washed three times with PBX, and incubated with 0.5 μg/ml DAPI at room temperature for 15 min. Samples were washed with PBX and mounted and imaged as described for FISH.

### Image processing

#### RNA FISH

Confocal images for RNA FISH were exported from Leica LAS-X. Brightness and contrast was adjusted in Image J-FIJI using these settings: probes for *Ars2* and *ZC3H18* in *w^shRNA^*, *Ars2^shRNA^*\*, and *ZC3H18^shRNA^* fly ovaries (minimum to maximum displayed values for DAPI channel was 0–100, *Ars2*-Qusar670 channel was 15–50, *ZC3H18*-Qusar570 channel was 10–100, and WGA-Alexa488 channel was 0–255), probes for *Rrp6* and *Dis3* in *w^shRNA^*, *Rrp6^shRNA^*, and *Dis3^shRNA^* fly ovaries (minimum to maximum displayed values for DAPI channel was 0 to 200, *Rrp6*-Qusar670 channel was 5–80, and *Dis3*-Qusar570 channel was 0 to 80), and probes for *IntS11* in *w^shRNA^* and *IntS11^shRNA^* fly ovaries (minimum to maximum displayed values for DAPI channel was 0–200 and *IntS11*-Qusar570 channel was 0–80). Individual grayscale channels used these setting: DAPI (200–255), *Ars2* (170–255), *ZC3H18* (100–255), *Rrp6* (150–255), *Dis3* (30–255), and *IntS11* (50–255). Image orientation was adjusted in Photoshop 24.4.0 (Adobe). Brightness of DAPI was set to the minimum value (−150) in Photoshop to avoid obscuring the transcript signal in the merged images. Z-overlayed images were generated using the average intensity of each stack.

#### Immunofluorescence

Confocal images for RNA FISH were exported from Leica LAS-X. Brightness and contrast was adjusted in Image J2-FIJI (v2.14) using these settings: minimum to maximum displayed values for DAPI channel (50–55), FLAG-CoraLite Plus 488 channel (10–100), Grk-CoraLite Plus 647 channel (10–100), and Lamin-CoraLite Plus 555 channel (20–255). Image orientation was adjusted in Photoshop. Images of germaria were 250 px × 250 px; for stages 5–7 they were 500 px × 500 px; for stages 8–10 they were 1000 px × 1000 px. Developmental timing was determined according to ref. (Jia et al., 2016).

#### RNA isolation

Fly ovaries were dissected into ice-cold PBS and washed twice with cold PBS, frozen in liquid nitrogen, and stored at −80°C. Ten volumes per ovary mass of mirVana Lysis/Binding buffer was added to frozen ovaries and the tissue disrupted using a motorized rotor-stator homogenizer (Thermo Fisher Scientific K7495400000) fitted with a plastic pestle (Thermo Fisher Scientific 2141364). Total RNA was purified using the mirVana kit (Life Technologies, #AM1561) according to the manufacturer’s instructions.

#### Total ovary RNA sequencing

mRNA sequencing libraries were constructed as previously described(Zhang et al., 2012) with modifications. ERCC spike-ins were added to 2 µg total RNA before library construction.(Baker et al., 2005; External RNA Controls Consortium, 2005) A pool of 186 fly rRNA-complementary oligonucleotides (0.05 μM each)(Adiconis et al., 2013; Morlan et al., 2012) was added to total RNA in hybridization buffer (10 mM Tris-HCl, pH 7.4, 20 mM NaCl), heated to 95°C, then cooled to 22°C at −0.1°C/sec. Thermostable RNase H (Lucigen, #H39500) was added and incubated at 45°C for 30 min in 50 mM Tris-Cl (pH 7.4), 100 mM NaCl, and 20 mM MgCl_2_. Next, Turbo DNase (Thermo Fisher #AM2238) was added (f.c. 0.08 U/μl) and the reaction incubated at 37°C for 20 min. RNAs >200 nt were purified using RNA Clean & Concentrator-5 (Zymo Research #R1016) according to the manufacturer’s instructions. RNA-seq libraries were sequenced using the Illumina NextSeq500 or 550 (strains expressing shRNA targeting *w, IntS11, IntS9, Rrp6, Dis3, ZC3H18, Ars2, and Prp19*) or NextSeq2000 (strains expressing shRNA targeting both *Dis3* and *Rrp6,* and strains expressing lhRNA targeting *RBM7, Mtr4, ZCCHC8*) using 75 + 75 nt, paired-end reads.

#### Oxford Nanopore Technologies (ONT) long-read sequencing of chromatin-associated, capped, non-polyadenylated RNA

Nascent transcripts were isolated as reported.(Khodor et al., 2011), with modifications. Ovaries from 3–5-day-old flies fed yeast for 2 days were homogenized with a Dounce homogenizer for 30 strokes with a tight type B pestle in 500 μl 15 mM HEPES-KOH, pH 7.6, 10 mM KCl, 5 mM magnesium acetate, 3 mM CaCl_2_, 300 mM sucrose, 0.1% (v/v) Triton X-100, 1 mM DTT, and the protease inhibitor cocktail described above. The lysate was strained through Miracloth (VWR # EM475855-1R) and 1 ml of 15 mM HEPES-KOH, pH 7,6, 10 mM potassium chloride, 5 mM magnesium acetate, 3 mM CaCl_2_, 1 M sucrose, 0.1% Triton X-100, 1 mM DTT, protease inhibitors was added. Lysate was centrifuged at 5,900 × *g* for 15 min at 4°C to pellet nuclei. Nuclei were resuspended in five volumes nuclear lysis buffer (10 mM HEPES-KOH, pH 7.6, 100 mM KCl, 0.1 mM EDTA, 10% (v/v) glycerol, 0.1 mM ZnCl_2_, 0.15 mM spermine, 0.5 mM spermidine, 10 mM NaF, 0.2 mM Na_3_VO_4_, 1 mM DTT, protease inhibitor, 40 U/µl RNasin Plus (Promega #N2615) and crushed 10 times with a plastic pestle. An equal volume of 2× NUN buffer (50 mM HEPES-KOH, pH 7.6, 600 mM NaCl, 2 M urea, 2% (v/v) NP-40, protease inhibitor) was added to the lysate dropwise and incubated on ice for 20 min. Nuclei were collected by centrifugation at 16,100 × *g* for 30 min at 4°C, and the nuclear pellet washed with a 1:1 solution of nuclear lysis buffer and 2× NUN buffer. RNA was purified using the mirVana RNA isolation kit according to the manufacturer’s instructions. Chromatin-associated RNA was treated with Quick calf intestinal alkaline phosphatase (New England Biolabs) in CutSmart buffer (50 mM potassium acetate, 20 mM Tris-acetate, 10 mM magnesium acetate, 100 µg/ml bovine serum albumin, pH 7.9) at 37°C for 30 min. Quick-CIP treated RNAs were purified with 1.8 vol RNAClean XP beads (Beckman Coulter), then incubated with 10 U/µl T4 polynucleotide kinase in T4 PNK reaction buffer (70 mM Tris-HCl, 10 mM magnesium chloride, 1 mM ATP, 5 mM DTT, pH 7.6) at 37°C for 30 min. RNA was purified with 1.8 vol RNAClean XP beads. RNA was incubated with 2.5 U Terminator exoribonuclease (Epicentre) in 20 µl buffer supplied by the manufacturer at 30°C for 1 h. The reaction was stopped by adding 5 mM EDTA. Poly(A)-tailed RNA (poly(A)^+^) or non-polyadenylated (poly(A)^−^) were separated using oligo (dT) magnetic beads (New England Biolabs S1419S). Briefly, an equal volume of binding buffer (20 mM Tris-HCl, pH 7.5, 1 M LiCl, 2 mM EDTA) was added to the RNA reaction mixture and incubated at 65°C for 2 min, then immediately chilled on ice for 2 min. Five μg oligo(dT)_25_ magnetic beads were washed three times with 200 μl binding buffer. Washed beads were added to the RNA mixture and incubated rotating at room temperature for 10 min. The supernatant, containing poly(A)^−^ RNA, was further purified with RNAClean XP beads (Beckman Coulter). Poly(A)^+^ RNA, bound to beads, was washed with 10 mM Tris-Cl, pH 7.5, 0.15 mM LiCl, and 1 mM EDTA, and eluted in water at 65°C for 2 min. Next, RNA was incubated with 1.25 µM single-stranded 3′ RNA adapter and 60 U RNA ligase 1 (NEB) in 20 µl ligation buffer (100 mM Tris-HCl, 50 mM magnesium chloride, 10 mM ATP, 100 mM DTT) at 16°C overnight. Ligated RNA was purified with 1.8 vol RNAClean XP beads. Next, cDNA was synthesized by incubating 1 μM reverse transcription primer (complementary to the 3′ adapter), 500 nM 5′ iso-template switch oligo, and 200 U Maxima H Minus Reverse Transcriptase (Thermo Fisher Scientific) at 50°C for 2 h, then heated at 85°C for 5 min (Table S9). To remove RNA template, 1 μl of RNase A/T1 mix (Thermo Fisher scientific) containing 2 mg/ml of RNase A and 5000 U/ml of RNase T1 was added to the reaction and incubated at 37°C for 10 min. cDNA libraries were amplified by PCR with Platinum SuperFi II DNA Polymerase (Thermo Fisher Scientific) as follows: 98°C for 30 sec, 14 cycles of 98°C for 10 sec, 65°C for 30 sec, 72°C for 10 min, then 72°C for 5 min. Amplified libraries were treated with 20 U exonuclease I (NEB) at 37°C for 15 min; the exonuclease was heat inactivated by 80°C for 5 min. cDNA libraries were purified using Ampure XP beads and libraries were constructed using the DNA ligation kit (Oxford Nanopore Technologies, SQK-109). Libraries were loaded onto R9.4.1 or R10.4.1 flow cells and sequenced using MinION (Table S10).

### Bioinformatic analyses

#### Total RNA sequencing

Barcodes were sorted, adapter sequences trimmed, and reads reformatted to extract UMIs. Alignment of RNA sequencing data to the *Drosophila melanogaster* genome (Dm6) was performed using piPipes 1.4.(Han et al., 2015) Reformatted reads were first aligned to rRNA using Bowtie 2 2.2.0.(Langmead and Salzberg, 2012) Unaligned unique reads were subsequently mapped to the Dm6 genome using STAR 2.3.1(Dobin et al., 2013), and multimapping reads removed using BEDTOOLS 2.30.0.(Quinlan and Hall, 2010)

#### ONT long-read sequencing of chromatin-associated, capped, non-polyadenylated RNA

Fast5 files were base-called using Guppy (Oxford Nanopore Technologies) and the resulting fastq files were aligned to the Dm6 genome using Minimap2 with default settings employed for cDNA ONT sequencing. Reads with both 5′ and 3′ adapter sequences were selected using Porechop.(Wick et al., 2017) Filtered FASTQ files were subsequently aligned to the Dm6 genome using Minimap2 with default settings employed for aligning cDNA ONT-Seq reads.(Li, 2018) Supplementary alignment-containing, multimapping and low-quality reads were subsequently discarded using in-house scripts.

#### Generation of genome browser tracks and metaplot analyses

Dm6-aligned reads were converted to per-million-scaled bigwig files using in-house scripts and visualized using the Integrative Genomics Viewer.(Robinson et al., 2011) Metaplots were generated by (1) converting Dm6-aligned ONT and Illumina-Seq reads to bigwig; (2) calculating the per-million scaled number of reads using DEEPTOOLS 3.5.1;(Ramírez et al., 2016), (3) normalizing values based on the average of the maximum values from the plotted genomic windows, and (4) using Lowess normalization for smoothing. Plots were generated using R 4.3.3.

#### Analysis of read-through transcription at snRNA loci using short-read RNA sequencing

Dm6-aligned short-read sequences were converted to bedGraph format and separately intersected with (1) the coordinates of Pol II-dependent non-intronic snRNAs (snRNALoc) and (2) 2000 bp regions directly downstream of the annotated snRNA transcription end site (DoG) using BEDTOOLS 2.30.0. The analysis was constrained to snRNAs lacking annotated genes in DoG, resulting in a total of 10 total loci. Normalized read-through was subsequently calculated as (DoG rpkm)/(snRNALoc rpkm) for each locus analyzed.

#### Comparison of snRNA length from long-read sequencing

Dm6-aligned ONT reads whose 5′ ends corresponded to the annotated transcription start sites of Pol II-dependent snRNAs were identified and their lengths recorded. Random resampling with replacement to determine 95% confidence intervals was performed using R 4.3.3.

#### Calculation of normalized PROMPT abundance from short-read RNA sequencing

Dm6-aligned Illumina-Seq reads were converted to bedGraph format and separately intersected with (1) the coordinates of the most upstream Dm6-annotated 5′ UTR for each gene and (2) PROMPT regions (genomic regions spanning 2000 bp upstream of the identified 5′ UTR on the opposite strand) using BEDTOOLS 2.30.0. 5′ UTR-PROMPT pairs harboring Dm6-annotated features within the defined PROMPT region were discarded, resulting in a total of 5,888 PROMPT loci with divergent 5′ UTRs. The gene-specific ratio of length-normalized reads overlapping PROMPT relative to length-normalized reads overlapping the divergent 5′ UTRs was calculated as (PROMPT read abundance/2000)/(length-normalized 5′ UTR read abundance). Regions with a calculated ratio >1 in the control were discarded to remove biases arising from incomplete genome annotation, resulting in 2,527 individual divergent PROMPT and 5′ UTR pairs.

#### Splicing efficiency

The coordinates of Dm6 ENSEMBL-annotated exon-exon junctions harboring ≥1 uniquely mapped reads in the control were extracted using default STAR 2.3.1 settings. CIGAR strings were parsed to identify split and unsplit (unspliced) alignments; junction coordinates identified by STAR were used as input for BEDTOOLS 2.30.0 to identify unsplit, unspliced alignments which each fully overlapped (1) regions defined as the 12 nt upstream and 3 nt downstream of the intron-exon (IE) boundaries or (2) regions defined as the 3 nt upstream and 12 nt downstream of the exon-intron (EI) boundaries. In parallel, junction coordinates identified by BEDTOOLS 2.30.0 and the STAR were used to identify individual split alignments that fully overlapped 7 nt directly upstream and downstream of exon-exon (EE) junctions, following the strategy detailed in ref.(Sánchez-Escabias et al., 2022). Splice sites with ≥1 read aligning to either the IE or EI boundary and ≥1 read at EE junctions in all genotypes were used (2,509 total junctions) to calculate a per-splice site metric of intron retention as (EI+IE)/(EI+IE+EE).

#### Statistical analyses

Statistical analyses were performed using base R 4.3.3 and related packages; BH-corrected statistics for post-hoc Dunn’s test were computed using the FSA package.(Ogle et al., 2025)

#### Identification and quantification of cryptic transcripts

GRO-seq data from three biological replicates of *Drosophila melanogaster* ovary tissue were merged to increase coverage and improve detection of actively transcribed regions.(Chang et al., 2019) Briefly, HOMER 4.11 was used to call peaks to identify putative transcription start site and transcriptional activity along the genome.(Heinz et al., 2010) Transcription start site and downstream loci were identified by filtering for enrichment scores >4 and >3, respectively. BEDTools 2.30.0, was used to perform strand-specific assignment of these transcriptionally active regions to Dm6 genomic regions lacking annotated loci.(Quinlan and Hall, 2010) Transcriptionally active regions overlapping or within 2,000 bp of Dm6-annotated loci were excluded, resulting in 1,014 cryptic transcripts from unannotated regions of the Dm6 genome. Reads overlapping the novel cryptic transcripts by at least 50% were extracted and used calculate the read length distribution.

---

**Table S3.** Related to S6A. Benjamini-Hochberg-corrected Mann-Whitney-Wilcoxon test *p*-values obtained from comparing the length distribution of reads aligning to each listed snRNA locus in the different germline shRNA-expressing ovaries compared to control.

| snRNA<br>locus | *w<sup>shRNA</sup>* control versus: |  |  |  |  |
| --- | --- | --- | --- | --- | --- |
|  | *Ars2<sup>shRNA</sup>* | *ZC3H19<sup>shRNA</sup>* | *Dis3<sup>shRNA</sup>* | *Rrp6<sup>shRNA</sup>* | *IntS11<sup>shRNA</sup>* |
| **U1:21D** | 1.0 (NS) | 1.0 (NS) | 0.94 (NS) | 1.0 (NS) | 0.0019 |
| **U1:82Eb** | 0.0011 | 0.18 (NS) | 0.041 | 0.061 | $3.6 \times 10^{-18}$ |
| **U1:95Ca** | 0.23 (NS) | 0.52 (NS) | 0.65 (NS) | 0.23 (NS) | $2.5 \times 10^{-17}$ |
| **U1:96Cb** | 0.0095 | 0.95 (NS) | 0.0095 | 0.95 (NS) | $4.2 \times 10^{-11}$ |
| **U1:95Cc** | 0.15 (NS) | 0.63 (NS) | 0.00060 | 0.63 (NS) | $7.2 \times 10^{-11}$ |
| **U2:14B** | 0.33 (NS) | 0.33 (NS) | 0.00030 | 0.73 (NS) | $5.4 \times 10^{-19}$ |
| **U4:39B** | 1.0 (NS) | 0.057 | 0.00024 | 1.0 (NS) | $9.0 \times 10^{-54}$ |
| **U11** | 0.72 (NS) | 0.72 (NS) | 0.32 (NS) | 0.72 (NS) | $4.3 \times 10^{-18}$ |
| **U12** | $1.8 \times 10^{-10}$ | $1.0 \times 10^{-18}$ | $9.4 \times 10^{-60}$ | $7.0 \times 10^{-7}$ | 0 |

**Table S4.** Related to S6A. Median ± SEM, estimated mean, lower and upper 95% confidence intervals (CI) obtained by sampling 100 times with replacement the length distributions for each snRNA locus in each strain expressing an shRNA in the female germline.

| snRNA locus | Median ± SEM | Est. Mean | Lower CI | Upper CI |
| --- | --- | --- | --- | --- |
| **U1:21D** | 334 ± 3 | 273 | 267 | 278 |
| **U1:82Eb** | 254 ± 3 | 308 | 301 | 314 |
| **U1:95Ca** | 339 ± 2 | 340 | 335 | 345 |
| **U1:95Cb** | 320 ± 2 | 311 | 307 | 315 |
| **U1:95Cc** | 215 ± 3 | 260 | 255 | 266 |
| **U2:14B** | 273 ± 3 | 333 | 326 | 339 |
| **U4:39B** | 147 ± 1 | 161 | 159 | 163 |
| **U11** | 271 ± 2 | 271 | 266 | 276 |
| **U12** | 245.0 ± 0.1 | 263.0 | 262.7 | 263.2 |

*IntS11<sup>shRNA</sup>*
| snRNA locus | Median ± SEM | Est. Mean | Lower CI | Upper CI |
| --- | --- | --- | --- | --- |
| **U1:21D** | 430 ± 8 | 554 | 538 | 571 |
| **U1:82Eb** | 640 ± 10 | 830 | 810 | 850 |
| **U1:95Ca** | 597 ± 7 | 662 | 648 | 677 |
| **U1:95Cb** | 480 ± 10 | 670 | 650 | 700 |
| **U1:95Cc** | 710 ± 10 | 840 | 820 | 870 |
| **U2:14B** | 845 ± 7 | 775 | 762 | 788 |
| **U4:39B** | 540 ± 5 | 662 | 652 | 672 |
| **U11** | 511 ± 7 | 555 | 540 | 569 |
| **U12** | 302.0 ± 0.6 | 439.2 | 438.0 | 440.3 |

**Ars2<sup>shRNA</sup>**
| snRNA locus | Median ± SEM | Est. Mean | Lower CI | Upper CI |
| --- | --- | --- | --- | --- |
| U1:21D | 363 ± 3 | 267 | 261 | 279 |
| U1:82Eb | 216 ± 3 | 235 | 230 | 240 |
| U1:95Ca | 355 ± 3 | 382 | 377 | 387 |
| U1:95Cb | 260 ± 4 | 277 | 270 | 284 |
| U1:95Cc | 165 ± 3 | 223 | 217 | 229 |
| U2:14B | 200 ± 3 | 278 | 272 | 284 |
| U4:39B | 147.0 ± 0.5 | 155 | 154. | 156. |
| U11 | 259 ± 3 | 266 | 260 | 272 |
| U12 | 246.0 ± 0.1 | 278.5 | 278.2 | 278.8 |

**ZC3H18<sup>shRNA</sup>**
| snRNA locus | Median ± SEM | Est. Mean | Lower CI | Upper CI |
| --- | --- | --- | --- | --- |
| U1:21D | 370 ± 3 | 312 | 306 | 318 |
| U1:82Eb | 241 ± 2 | 259 | 254 | 263 |
| U1:95Ca | 366 ± 5 | 415 | 405 | 425 |
| U1:95Cb | 304 ± 2 | 305 | 301 | 310 |
| U1:95Cc | 168 ± 3 | 222 | 216 | 228 |
| U2:14B | 380 ± 4 | 419 | 410 | 427 |
| U4:39B | 146 ± 1 | 157 | 155 | 159 |
| U11 | 275 ± 3 | 310 | 304 | 316 |
| U12 | 244.0 ± 0.1 | 266.6 | 266.4 | 266.9 |

***Dis3*<sup>shRNA</sup>**
| snRNA locus | Median ± SEM | Est. Mean | Lower CI | Upper CI |
| --- | --- | --- | --- | --- |
| U1:21D | 274 ± 2 | 255 | 250 | 259 |
| U1:82Eb | 232 ± 2 | 245 | 241 | 249 |
| U1:95Ca | 300 ± 3 | 355 | 348 | 362 |
| U1:95Cb | 265 ± 3 | 272 | 266 | 277 |
| U1:95Cc | 161 ± 2 | 180 | 176 | 1832 |
| U2:14B | 197 ± 2 | 245 | 241 | 249 |
| U4:39B | 145.0 ± 0.3 | 146.5 | 145.9 | 147.1 |
| U11 | 249 ± 2 | 253 | 249 | 257 |
| U12 | 243.0 ± 0.1 | 256.0 | 255.8 | 256.2 |

***Rrp6*<sup>shRNA</sup>**
| snRNA locus | Median ± SEM | Est. Mean | Lower CI | Upper CI |
| --- | --- | --- | --- | --- |
| U1:21D | 447 ± 5 | 301 | 292 | 310 |
| U1:82Eb | 270 ± 8 | 461 | 445 | 476 |
| U1:95Ca | 355 ± 7 | 485 | 471 | 499 |
| U1:95Cb | 311 ± 2 | 294 | 291 | 297 |
| U1:95Cc | 404 ± 7 | 429 | 414 | 443 |
| U2:14B | 200 ± 4 | 338 | 331 | 346 |
| U4:39B | 147 ± 3 | 231 | 225 | 237 |
| U11 | 265 ± 4 | 300 | 291 | 309 |
| U12 | 246.0 ± 0.1 | 276.5 | 276.2 | 276.8 |

**Table S5.** Related to Figure S6B and S6D. Spearman correlation (ρ) between short-read sequencing data sets.

| **Trial 1 vs. Trial 2** | **$\rho$** |
| --- | --- |
| *w*<sup>shRNA</sup> | 0.95 |
| *IntS11*<sup>shRNA</sup> | 0.95 |
| *IntS9*<sup>shRNA</sup> | 0.95 |
| *Rrp6*<sup>shRNA</sup> | 0.94 |
| *Dis3*<sup>shRNA</sup> | 0.95 |
| *ZC3H18*<sup>shRNA</sup> | 0.94 |
| *Ars2*<sup>shRNA</sup> | 0.95 |
| *Dis3, Rrp6*<sup>shRNA</sup> | 0.95 |
| *Mtr4*<sup>shRNA</sup> | 0.93 |
| *Rbm7*<sup>shRNA</sup> | 0.93 |
| *ZCCHC8*<sup>shRNA</sup> | 0.94 |

**Table S6.**
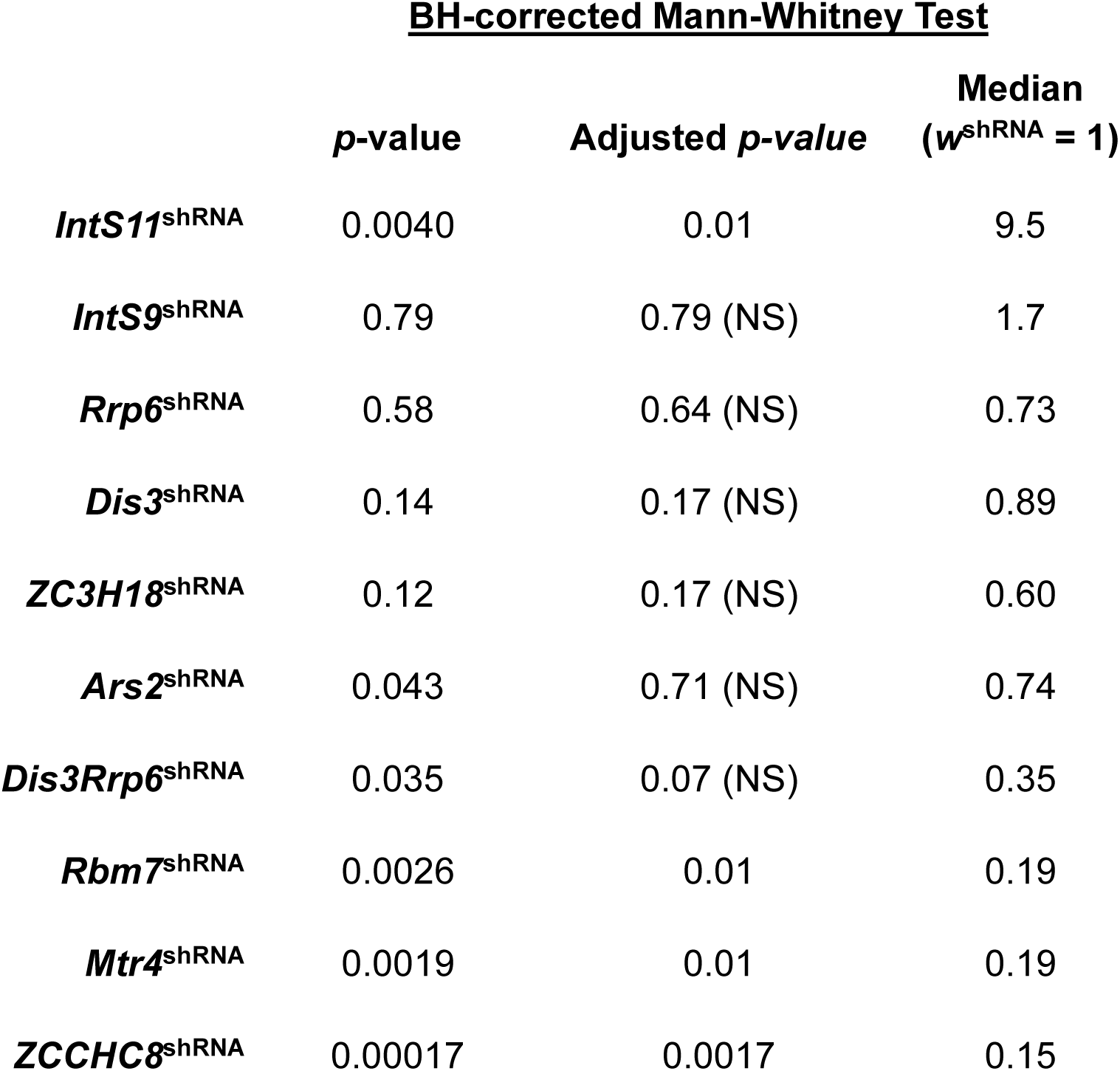
Related to Figure S6D. Benjamini-Hochberg-corrected Mann-Whitney-Wilcoxon test *p-*value and median relative to control for each strain expressing an shRNA in the female germline.

## SUPPLEMENTAL INFORMATION

**Figure S1, related to Figure 3.**
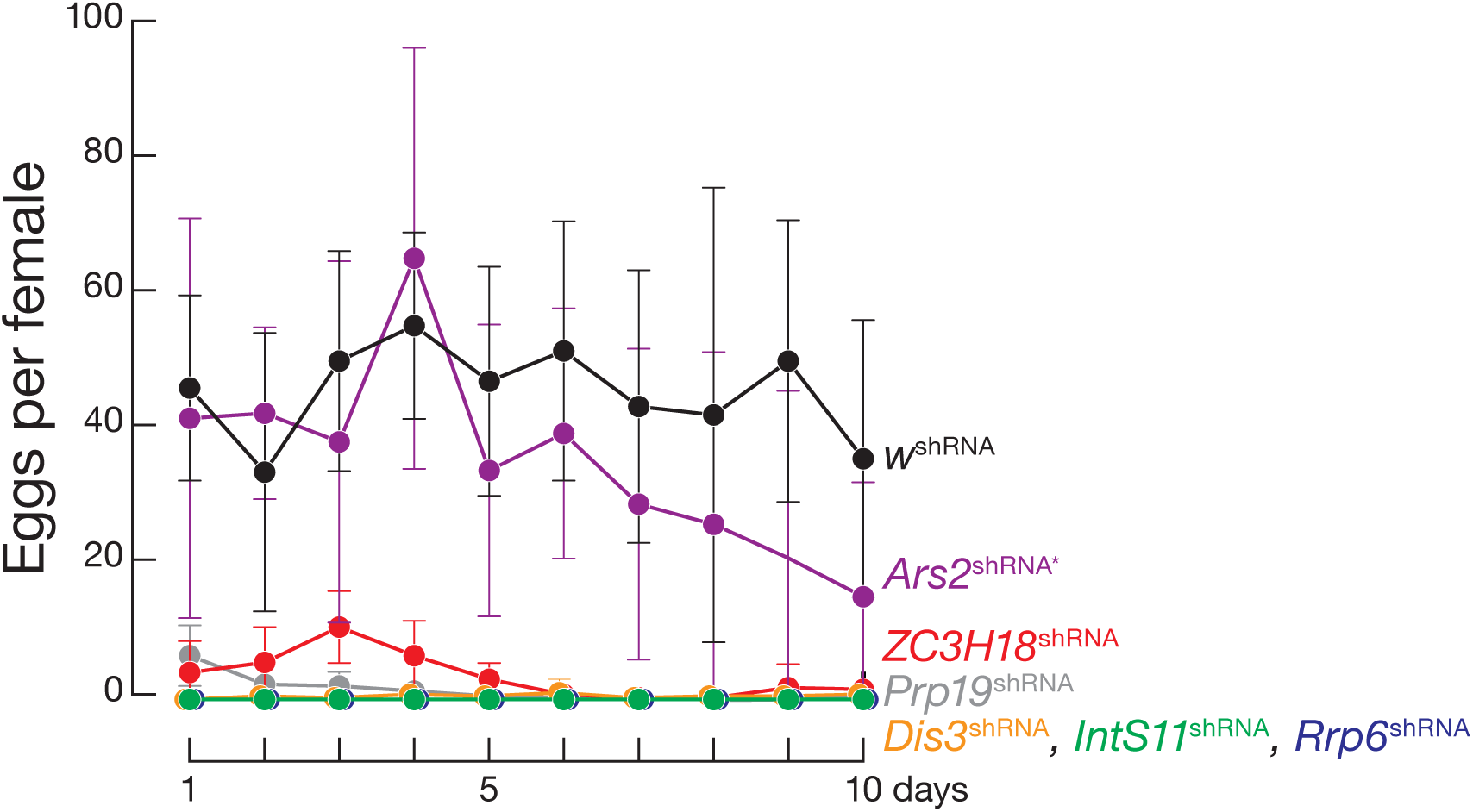
Components of the Integrator, NEXT, and nuclear exosome are required for *Drosophila* female fertility. Eggs laid per female per day. Four females were tested for each shRNA target gene. Data are mean ± SD. Females expressing the shRNA targeting *Ars2* were heterozygous for the loss-of-function allele *Ars2^R12fsX^* (*Ars2*^shRNA*^).

**Figure S2, related to Figure 3.**
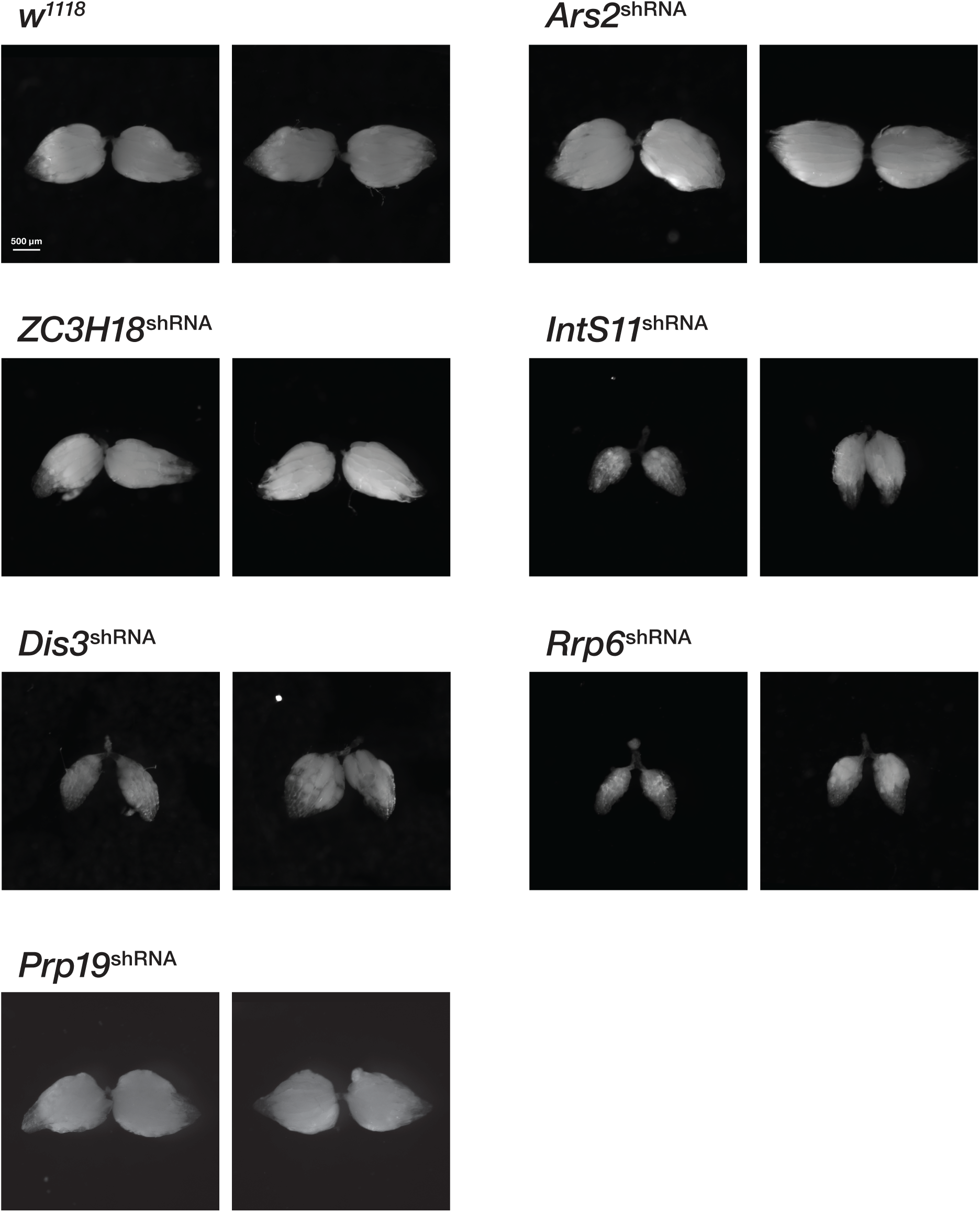
Depletion of Integrator, NEXT, and nuclear exosome components arrests oogenesis before stage 14. Bright-field images of ovaries expressing in the germ line an shRNA targeting the indicated gene were taken of the dissected ovaries on the fifth day.

**Figure S3, related to Figure 4.**
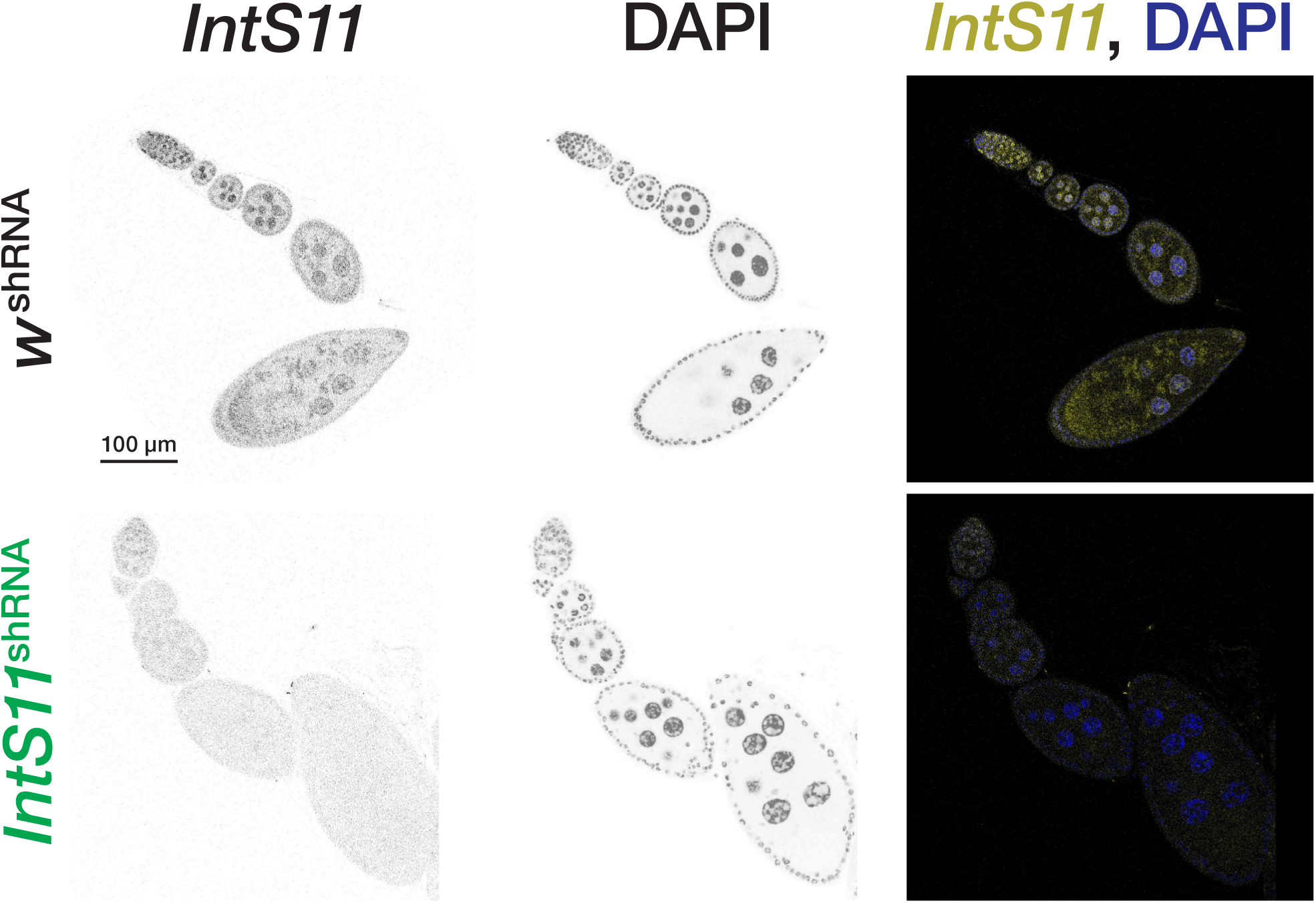
*IntS11* mRNA can be detected until oogenesis stage 10. DNA was detected with DAPI (blue); *IntS11* was detected by FISH (yellow). Females expressing the shRNA targeting *Ars2* were heterozygous for the loss-of-function allele *Ars2^R12fsX^* (*Ars2^shRNA^*\*).

**Figure S4, related to Figure 4.**
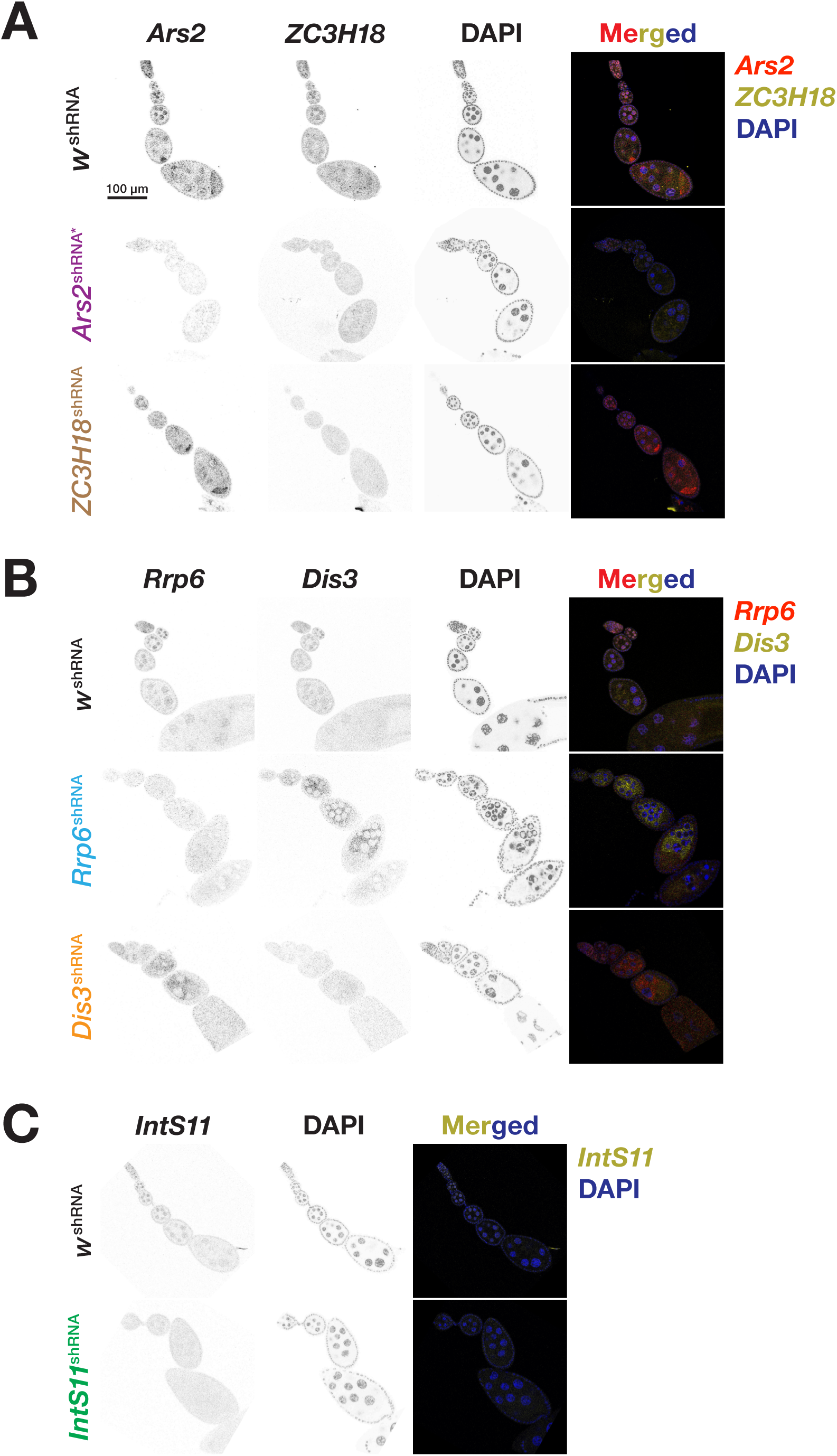
Additional RNA FISH images for ovaries expressing in the germ line shRNAs targeting *Ars2*, *ZC3H18*, *Rrp6, Dis3*, or *IntS11*. (A–C) RNA localization of Integrator, NEXT, or nuclear exosome components was detected by FISH using a pool of 48 Stellaris probes complementary to the transcript of the gene at the top of each column. Ovaries were dissected from three-days-old female flies and imaged using an SP-8 Leica confocal microscope. Nuclei were stained with DAPI (blue). A series of 20–25 images were collected to produce Z-stacked images.

**Figure S5, related to Figure 5.**
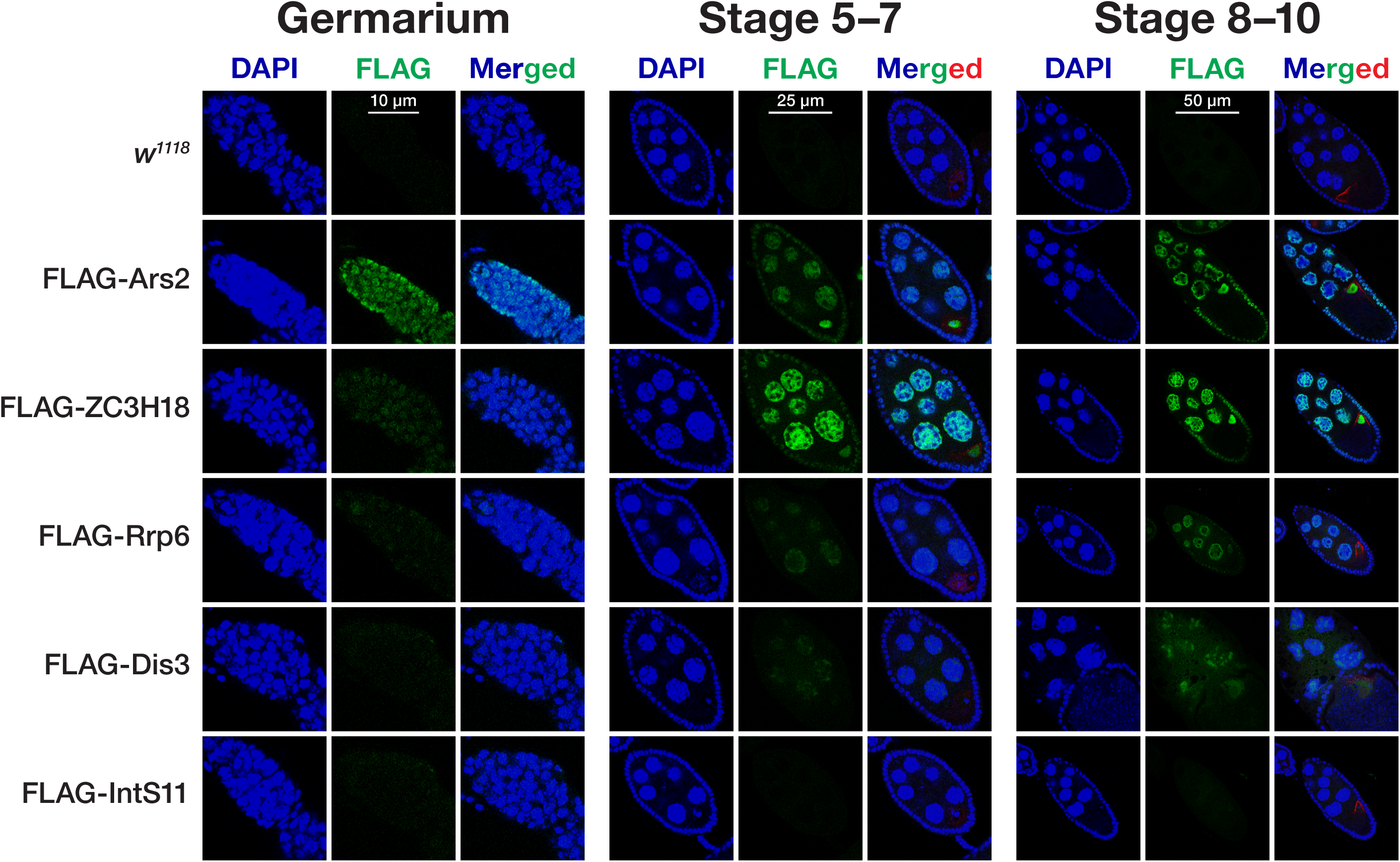
Expression and localization of epitope-tagged proteins during *Drosophila* oogenesis. Localization of 3XFLAG-epitope tagged Ars2, ZC3H18, Rrp6, Dis3, or IntS11 protein in the germarium, pre-vitellogenic stages, and vitellogenic stages of oogenesis was detected using anti-FLAG M2 conjugated to CoraLite Plus 488. Gurken was detected using anti-Gurken antibody conjugated to CoraLite Plus 647. Nuclei were stained with DAPI (blue). *w^1118^* serves as a negative control.

**Figure S6, related to Figure 6.**
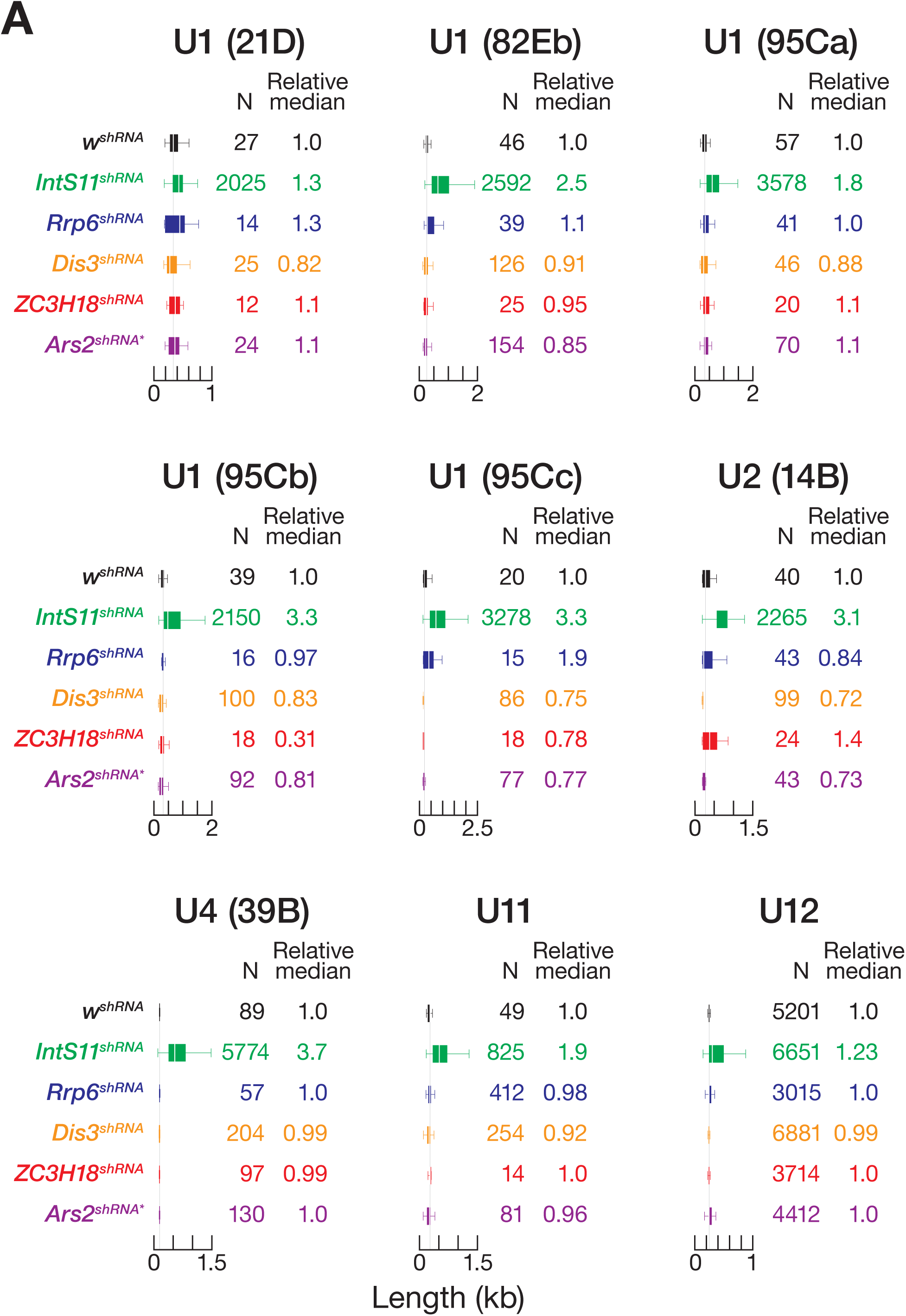
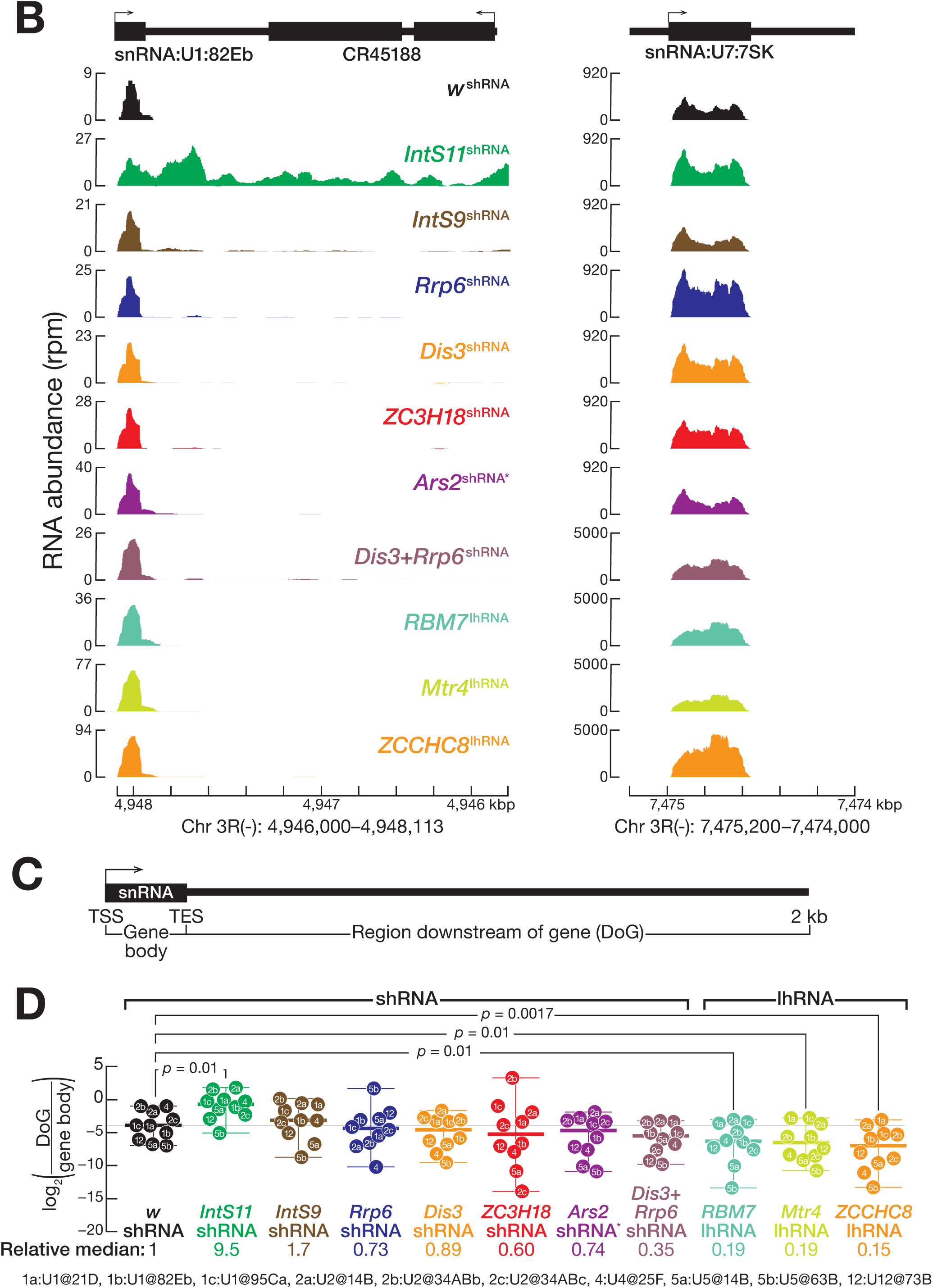
Total RNA abundance for Pol II-dependent snRNA, Pol III-dependent 7SK snRNA, and for protein-coding loci. (A) Median length of long-read sequences whose 5′ end begins at the annotated transcription start site of Pol II-dependent snRNAs. Boxplots depict the inter-quartile ranges, median, maximum and minimum values after excluding outliers. N, number of reads. (B) The abundance of RNA mapping to the U1 snRNA locus at 82Eb or the 7SK locus, measured using short-read sequencing of ovary RNA, for flies expressing in the female germ line an shRNA or lhRNA targeting the gene indicated. *RBM7*, *Mtr4*, and *ZCCHC8* were depleted using long-hairpin RNA instead of an shRNA. (C) Strategy used to calculate the abundance of downstream of gene (DoG) transcripts. (D) The abundance of DoG RNA, relative to gene body transcripts was measured using short-read sequencing for ten Pol II-dependent snRNA loci. The loci were selected because they do not overlap with annotated protein-coding or long-non-coding genes. Plots show median, maximum, and minimum values after excluding outliers. Code names inside the markers indicate the specific snRNA identity and chromosomal location (e.g., 1a indicates U1 snRNA at 21D, abbreviated U1@21D), as listed at the bottom of the panel. BH-corrected Mann-Whitney Wilcoxon *p-*values compared to control are shown. *RBM7*, *Mtr4*, and *ZCCHC8* were depleted using long-hairpin RNA instead of an shRNA.

**Figure S7, related to Figure 7.**
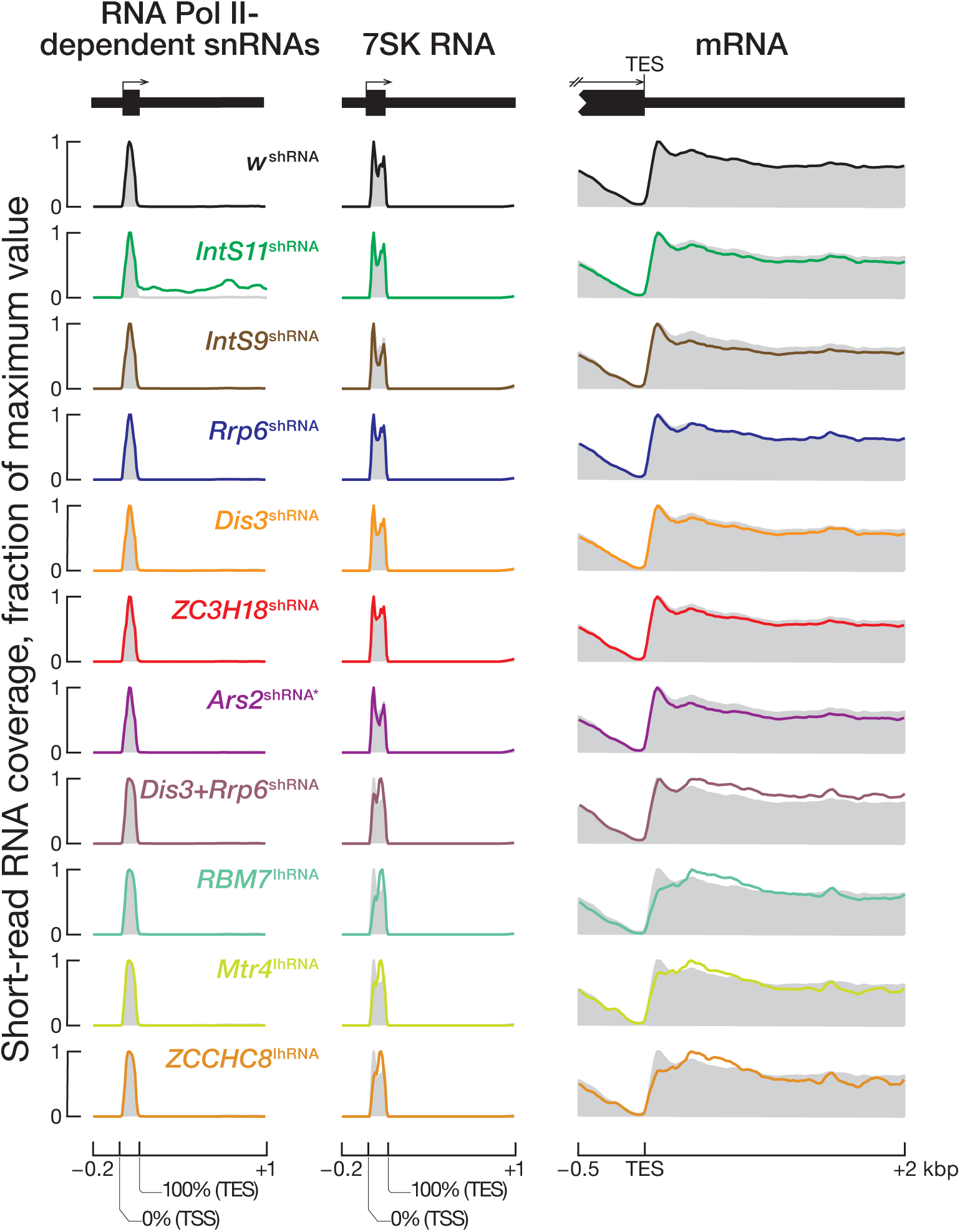
Metaplot analyses of the abundance of RNA mapping to Pol II-dependent snRNA, Pol III-dependent 7SK RNA, or protein coding loci as measured by short-read sequencing of total ovary RNA. Metaplot analysis of short-read sequencing data normalized to sequencing depth for 25 annotated Pol II-dependent snRNA loci (left), the Pol III-dependent 7SK locus, and 13,963 protein-coding genes. Gray shadows depict the abundance in the *w^shRNA^* control. RBM7, Mtr4, and ZCCHC8 were depleted using long-hairpin RNA instead of an shRNA.

## References

Adám, G., Gausz, J., Noselli, S., Kurucz, E., Andó, I., and Udvardy, A. (2004). Tissue- and developmental stage-specific changes in the subcellular localization of the 26S proteasome in the ovary of *Drosophila melanogaster*. Gene Expr Patterns 4, 329–333.

Adiconis, X., Borges-Rivera, D., Satija, R., Deluca, D. S., Busby, M. A., Berlin, A. M., Sivachenko, A., Thompson, D. A., Wysoker, A., Fennell, T., Gnirke, A., Pochet, N., Regev, A., and Levin, J. Z. (2013). Comparative analysis of RNA sequencing methods for degraded or low-input samples. Nat Methods

Albrecht, T. R., Shevtsov, S. P., Wu, Y., Mascibroda, L. G., Peart, N. J., Huang, K. L., Sawyer, I. A., Tong, L., Dundr, M., and Wagner, E. J. (2018). Integrator subunit 4 is a ‘Symplekin-like’ scaffold that associates with INTS9/11 to form the Integrator cleavage module. Nucleic Acids Res 46, 4241–4255.

Albrecht, T. R., and Wagner, E. J. (2012). snRNA 3’ end formation requires heterodimeric association of integrator subunits. Mol Cell Biol 32, 1112–1123.

Allmang, C., Mitchell, P., Petfalski, E., and Tollervey, D. (2000). Degradation of ribosomal RNA precursors by the exosome. Nucleic Acids Res 28, 1684–1691.

Allmang, C., Petfalski, E., Podtelejnikov, A., Mann, M., Tollervey, D., and Mitchell, P. (1999). The yeast exosome and human PM-Scl are related complexes of 3’ --> 5’ exonucleases. Genes Dev 13, 2148–2158.

Andersen, P. R., Domanski, M., Kristiansen, M. S., Storvall, H., Ntini, E., Verheggen, C., Schein, A., Bunkenborg, J., Poser, I., Hallais, M., Sandberg, R., Hyman, A., LaCava, J., Rout, M. P., Andersen, J. S., Bertrand, E., and Jensen, T. H. (2013). The human cap-binding complex is functionally connected to the nuclear RNA exosome. Nat Struct Mol Biol 20, 1367–1376.

Arkinson, C., Dong, K. C., Gee, C. L., and Martin, A. (2025). Mechanisms and regulation of substrate degradation by the 26S proteasome. Nat Rev Mol Cell Biol 26, 104–122.

Azuma, N., Yokoi, T., Tanaka, T., Matsuzaka, E., Saida, Y., Nishina, S., Terao, M., Takada, S., Fukami, M., Okamura, K., Maehara, K., Yamasaki, T., Hirayama, J., Nishina, H., Handa, H., and Yamaguchi, Y. (2023). Integrator complex subunit 15 controls mRNA splicing and is critical for eye development. Hum Mol Genet 32, 2032–2045.

Baillat, D., Hakimi, M. A., Naar, A. M., Shilatifard, A., Cooch, N., and Shiekhattar, R. (2005). Integrator, a multiprotein mediator of small nuclear RNA processing, associates with the C-terminal repeat of RNA polymerase II. Cell 123, 265–276.

Baker, S. C., Bauer, S. R., Beyer, R. P., Brenton, J. D., Bromley, B., Burrill, J., Causton, H., Conley, M. P., Elespuru, R., Fero, M., Foy, C., Fuscoe, J., Gao, X., Gerhold, D. L., Gilles, P., Goodsaid, F., Guo, X., Hackett, J., Hockett, R. D., Ikonomi, P., Irizarry, R. A., Kawasaki, E. S., Kaysser-Kranich, T., Kerr, K., Kiser, G., Koch, W. H., Lee, K. Y., Liu, C., Liu, Z. L., Lucas, A., Manohar, C. F., Miyada, G., Modrusan, Z., Parkes, H., Puri, R. K., Reid, L., Ryder, T. B., Salit, M., Samaha, R. R., Scherf, U., Sendera, T. J., Setterquist, R. A., Shi, L., Shippy, R., Soriano, J. V., Wagar, E. A., Warrington, J. A., Williams, M., Wilmer, F., Wilson, M., Wolber, P. K., Wu, X., Zadro, R., and External RNA Controls Consortium (2005). The External RNA Controls Consortium: a progress report. Nat Methods 2, 731–734.

Barbieri, E., Trizzino, M., Welsh, S. A., Owens, T. A., Calabretta, B., Carroll, M., Sarma, K., and Gardini, A. (2018). Targeted Enhancer Activation by a Subunit of the Integrator Complex. Mol Cell 71, 103–116.e7.

Bard, J. A. M., Goodall, E. A., Greene, E. R., Jonsson, E., Dong, K. C., and Martin, A. (2018). Structure and Function of the 26S Proteasome. Annu Rev Biochem 87, 697–724.

Barra, J., Gaidosh, G. S., Blumenthal, E., Beckedorff, F., Tayari, M. M., Kirstein, N., Karakach, T. K., Jensen, T. H., Impens, F., Gevaert, K., Leucci, E., Shiekhattar, R., and Marine, J. C. (2020). Integrator restrains paraspeckles assembly by promoting isoform switching of the lncRNA NEAT1. Sci Adv 6, eaaz9072.

Beckedorff, F., Blumenthal, E., daSilva, L. F., Aoi, Y., Cingaram, P. R., Yue, J., Zhang, A., Dokaneheifard, S., Valencia, M. G., Gaidosh, G., Shilatifard, A., and Shiekhattar, R. (2020). The Human Integrator Complex Facilitates Transcriptional Elongation by Endonucleolytic Cleavage of Nascent Transcripts. Cell Rep 32, 107917.

Chan, S.-P., Kao, D.-I., Tsai, W.-Y., and Cheng, S.-C. (2003). The Prp19p-associated complex in spliceosome activation. Science 302, 279–282.

Chang, T. H., Mattei, E., Gainetdinov, I., Colpan, C., Weng, Z., and Zamore, P. D. (2019). Maelstrom Represses Canonical Polymerase II Transcription within Bi-directional piRNA Clusters in *Drosophila melanogaster*. Mol Cell 73, 291–303.e6.

Chen, J., Ezzeddine, N., Waltenspiel, B., Albrecht, T. R., Warren, W. D., Marzluff, W. F., and Wagner, E. J. (2012). An RNAi screen identifies additional members of the *Drosophila* Integrator complex and a requirement for cyclin C/Cdk8 in snRNA 3’-end formation. RNA 18, 2148–2156.

Chiu, A. C., Suzuki, H. I., Wu, X., Mahat, D. B., Kriz, A. J., and Sharp, P. A. (2018). Transcriptional Pause Sites Delineate Stable Nucleosome-Associated Premature Polyadenylation Suppressed by U1 snRNP. Mol Cell 69, 648–663.e7.

Cho, U. S., and Xu, W. (2007). Crystal structure of a protein phosphatase 2A heterotrimeric holoenzyme. Nature 445, 53–57.

Cihlarova, Z., Kubovciak, J., Sobol, M., Krejcikova, K., Sachova, J., Kolar, M., Stanek, D., Barinka, C., Yoon, G., Caldecott, K. W., and Hanzlikova, H. (2022). BRAT1 links Integrator and defective RNA processing with neurodegeneration. Nat Commun 13, 5026.

Cortazar, M. A., Sheridan, R. M., Erickson, B., Fong, N., Glover-Cutter, K., Brannan, K., and Bentley, D. L. (2019). Control of RNA Pol II Speed by PNUTS-PP1 and Spt5 Dephosphorylation Facilitates Termination by a “Sitting Duck Torpedo” Mechanism. Mol Cell 76, 896–908.e4.

Cox, J., and Mann, M. (2008). MaxQuant enables high peptide identification rates, individualized p.p.b.-range mass accuracies and proteome-wide protein quantification. Nat Biotechnol 26, 1367–1372.

Cox, J., Neuhauser, N., Michalski, A., Scheltema, R. A., Olsen, J. V., and Mann, M. (2011). Andromeda: a peptide search engine integrated into the MaxQuant environment. J Proteome Res 10, 1794–1805.

Coy, S., Volanakis, A., Shah, S., and Vasiljeva, L. (2013). The Sm complex is required for the processing of non-coding RNAs by the exosome. PLoS One 8, e65606.

Dasilva, L. F., Blumenthal, E., Beckedorff, F., Cingaram, P. R., Gomes Dos Santos, H., Edupuganti, R. R., Zhang, A., Dokaneheifard, S., Aoi, Y., Yue, J., Kirstein, N., Tayari, M. M., Shilatifard, A., and Shiekhattar, R. (2021). Integrator enforces the fidelity of transcriptional termination at protein-coding genes. Sci Adv 7, eabe3393.

Davidson, L., Francis, L., Cordiner, R. A., Eaton, J. D., Estell, C., Macias, S., Cáceres, J. F., and West, S. (2019). Rapid Depletion of DIS3, EXOSC10, or XRN2 Reveals the Immediate Impact of Exoribonucleolysis on Nuclear RNA Metabolism and Transcriptional Control. Cell Rep 26, 2779–2791.e5.

Ogle, D. H., Doll, J. C., Wheeler, A. P., and Dinno, A. (2025). FSA: Simple Fisheries Stock Assessment Methods.

Dobin, A., Davis, C. A., Schlesinger, F., Drenkow, J., Zaleski, C., Jha, S., Batut, P., Chaisson, M., and Gingeras, T. R. (2013). STAR: ultrafast universal RNA-seq aligner. Bioinformatics 29, 15–21.

Dokaneheifard, S., Gomes Dos Santos, H., Guiselle Valencia, M., Arigela, H., Edupuganti, R. R., and Shiekhattar, R. (2024). Neuronal differentiation requires BRAT1 complex to remove REST from chromatin. Proc Natl Acad Sci U S A 121, e2318740121.

Dominski, Z., Yang, X. C., and Marzluff, W. F. (2005). The polyadenylation factor CPSF-73 is involved in histone-pre-mRNA processing. Cell 123, 37–48.

Dubiez, E., Pellegrini, E., Finderup Brask, M., Garland, W., Foucher, A. E., Huard, K., Heick Jensen, T., Cusack, S., and Kadlec, J. (2024). Structural basis for competitive binding of productive and degradative co-transcriptional effectors to the nuclear cap-binding complex. Cell Rep 43, 113639.

Egloff, S., Szczepaniak, S. A., Dienstbier, M., Taylor, A., Knight, S., and Murphy, S. (2010). The integrator complex recognizes a new double mark on the RNA polymerase II carboxyl-terminal domain. J Biol Chem 285, 20564–20569.

Egloff, S., Zaborowska, J., Laitem, C., Kiss, T., and Murphy, S. (2012). Ser7 phosphorylation of the CTD recruits the RPAP2 Ser5 phosphatase to snRNA genes. Mol Cell 45, 111–122.

ElMaghraby, M. F., Tirian, L., Senti, K. A., Meixner, K., and Brennecke, J. (2022). A genetic toolkit for studying transposon control in the *Drosophila melanogaster* ovary. Genetics 220, iyab179.

Elrod, N. D., Henriques, T., Huang, K. L., Tatomer, D. C., Wilusz, J. E., Wagner, E. J., and Adelman, K. (2019). The Integrator Complex Attenuates Promoter-Proximal Transcription at Protein-Coding Genes. Mol Cell 76, 738–752.e7.

Evans, R., O’Neill, M., Pritzel, A., Antropova, N., Senior, A., Green, T., Žídek, A., Bates, R., Blackwell, S., Yim, J., Ronneberger, O., Bodenstein, S., Zielinski, M., Bridgland, A., Potapenko, A., Cowie, A., Tunyasuvunakool, K., Jain, R., Clancy, E., Kohli, P., Jumper, J., and Hassabis, D. (2021). Protein complex prediction with AlphaFold-Multimer. bioRxiv 1846.

External RNA Controls Consortium (2005). Proposed methods for testing and selecting the ERCC external RNA controls. BMC Genomics 6, 150.

Falk, S., Finogenova, K., Melko, M., Benda, C., Lykke-Andersen, S., Jensen, T. H., and Conti, E. (2016). Structure of the RBM7-ZCCHC8 core of the NEXT complex reveals connections to splicing factors. Nat Commun 7, 13573.

Fang, Y., and Spector, D. L. (2007). Identification of nuclear dicing bodies containing proteins for microRNA biogenesis in living Arabidopsis plants. Curr Biol 17, 818–823.

Fianu, I., Chen, Y., Dienemann, C., Dybkov, O., Linden, A., Urlaub, H., and Cramer, P. (2021). Structural basis of Integrator-mediated transcription regulation. Science 374, 883–887.

Fianu, I., Ochmann, M., Walshe, J. L., Dybkov, O., Cruz, J. N., Urlaub, H., and Cramer, P. (2024). Structural basis of Integrator-dependent RNA polymerase II termination. Nature 629, 219–227.

Ge, D. T., Tipping, C., Brodsky, M. H., and Zamore, P. D. (2016). Rapid Screening for CRISPR-Directed Editing of the *Drosophila* Genome Using white Coconversion. G3 (Bethesda) 6, 3197–3206.

Gerlach, P., Garland, W., Lingaraju, M., Salerno-Kochan, A., Bonneau, F., Basquin, J., Jensen, T. H., and Conti, E. (2022). Structure and regulation of the nuclear exosome targeting complex guides RNA substrates to the exosome. Mol Cell 82, 2505–2518.e7.

Gómez-Orte, E., Sáenz-Narciso, B., Zheleva, A., Ezcurra, B., de Toro, M., López, R., Gastaca, I., Nilsen, H., Sacristán, M. P., Schnabel, R., and Cabello, J. (2019). Disruption of the *Caenorhabditis elegans* Integrator complex triggers a non-conventional transcriptional mechanism beyond snRNA genes. PLoS Genet 15, e1007981.

Goodrich, J. S., Clouse, K. N., and Schüpbach, T. (2004). Hrb27C, Sqd and Otu cooperatively regulate gurken RNA localization and mediate nurse cell chromosome dispersion in *Drosophila* oogenesis. Development 131, 1949–1958.

Graham, A. C., Kiss, D. L., and Andrulis, E. D. (2006). Differential distribution of exosome subunits at the nuclear lamina and in cytoplasmic foci. Mol Biol Cell 17, 1399–1409.

Gruber, J. J., Olejniczak, S. H., Yong, J., La Rocca, G., Dreyfuss, G., and Thompson, C. B. (2012). Ars2 promotes proper replication-dependent histone mRNA 3’ end formation. Mol Cell 45, 87–98.

Gruber, J. J., Zatechka, D. S., Sabin, L. R., Yong, J., Lum, J. J., Kong, M., Zong, W. X., Zhang, Z., Lau, C. K., Rawlings, J., Cherry, S., Ihle, J. N., Dreyfuss, G., and Thompson, C. B. (2009). Ars2 links the nuclear cap-binding complex to RNA interference and cell proliferation. Cell 138, 328–339.

Hallais, M., Pontvianne, F., Andersen, P. R., Clerici, M., Lener, D., Benbahouche, N. H., Gostan, T., Vandermoere, F., Robert, M. C., Cusack, S., Verheggen, C., Jensen, T. H., and Bertrand, E. (2013). CBC-ARS2 stimulates 3’-end maturation of multiple RNA families and favors cap-proximal processing. Nat Struct Mol Biol 20, 1358–1366.

Han, B. W., Wang, W., Zamore, P. D., and Weng, Z. (2015). piPipes: a set of pipelines for piRNA and transposon analysis via small RNA-seq, RNA-seq, degradome- and CAGE-seq, ChIP-seq and genomic DNA sequencing. Bioinformatics 31, 593–595.

Heinz, S., Benner, C., Spann, N., Bertolino, E., Lin, Y. C., Laslo, P., Cheng, J. X., Murre, C., Singh, H., and Glass, C. K. (2010). Simple combinations of lineage-determining transcription factors prime cis-regulatory elements required for macrophage and B cell identities. Mol Cell 38, 576–589.

Hocine, S., Singer, R. H., and Grünwald, D. (2010). RNA processing and export. Cold Spring Harb Perspect Biol 2, a000752.

Houseley, J., LaCava, J., and Tollervey, D. (2006). RNA-quality control by the exosome. Nat Rev Mol Cell Biol 7, 529–539.

Hrossova, D., Sikorsky, T., Potesil, D., Bartosovic, M., Pasulka, J., Zdrahal, Z., Stefl, R., and Vanacova, S. (2015). RBM7 subunit of the NEXT complex binds U-rich sequences and targets 3’-end extended forms of snRNAs. Nucleic Acids Res 43, 4236–4248.

Hu, Y., Flockhart, I., Vinayagam, A., Bergwitz, C., Berger, B., Perrimon, N., and Mohr, S. E. (2011). An integrative approach to ortholog prediction for disease-focused and other functional studies. BMC Bioinformatics 12, 357.

Huang, K. L., Jee, D., Stein, C. B., Elrod, N. D., Henriques, T., Mascibroda, L. G., Baillat, D., Russell, W. K., Adelman, K., and Wagner, E. J. (2020). Integrator Recruits Protein Phosphatase 2A to Prevent Pause Release and Facilitate Transcription Termination. Mol Cell 80, 345–358.e9.

Jambor, H., Surendranath, V., Kalinka, A. T., Mejstrik, P., Saalfeld, S., and Tomancak, P. (2015). Systematic imaging reveals features and changing localization of mRNAs in Drosophila development. Elife 4, e05003.

Janssens, V., and Goris, J. (2001). Protein phosphatase 2A: a highly regulated family of serine/threonine phosphatases implicated in cell growth and signalling. Biochem J 353, 417–439.

Jia, D., Xu, Q., Xie, Q., Mio, W., and Deng, W.-M. (2016). Automatic stage identification of Drosophila egg chamber based on DAPI images. Sci Rep 6, 18850.

Jumper, J., Evans, R., Pritzel, A., Green, T., Figurnov, M., Ronneberger, O., Tunyasuvunakool, K., Bates, R., Žídek, A., Potapenko, A., Bridgland, A., Meyer, C., Kohl, S. A. A., Ballard, A. J., Cowie, A., Romera-Paredes, B., Nikolov, S., Jain, R., Adler, J., Back, T., Petersen, S., Reiman, D., Clancy, E., Zielinski, M., Steinegger, M., Pacholska, M., Berghammer, T., Bodenstein, S., Silver, D., Vinyals, O., Senior, A. W., Kavukcuoglu, K., Kohli, P., and Hassabis, D. (2021). Highly accurate protein structure prediction with AlphaFold. Nature 596, 583–589.

Jurica, M. S., and Moore, M. J. (2003). Pre-mRNA splicing: awash in a sea of proteins. Mol Cell 12, 5–14.

Khodor, Y. L., Rodriguez, J., Abruzzi, K. C., Tang, C. H., Marr, M. T., and Rosbash, M. (2011). Nascent-seq indicates widespread cotranscriptional pre-mRNA splicing in *Drosophila*. Genes Dev 25, 2502–2512.

Kilchert, C., Wittmann, S., and Vasiljeva, L. (2016). The regulation and functions of the nuclear RNA exosome complex. Nat Rev Mol Cell Biol 17, 227–239.

Köhler, A., and Hurt, E. (2007). Exporting RNA from the nucleus to the cytoplasm. Nat Rev Mol Cell Biol 8, 761–773.

Lai, F., Gardini, A., Zhang, A., and Shiekhattar, R. (2015). Integrator mediates the biogenesis of enhancer RNAs. Nature 525, 399–403.

Langmead, B., and Salzberg, S. L. (2012). Fast gapped-read alignment with Bowtie 2. Nat Methods 9, 357–359.

Lei, E. P., and Silver, P. A. (2002). Protein and RNA export from the nucleus. Dev Cell 2, 261–272.

Lex, A., Gehlenborg, N., Strobelt, H., Vuillemot, R., and Pfister, H. (2014). UpSet: Visualization of Intersecting Sets. IEEE Trans Vis Comput Graph 20, 1983–1992.

Li, C., Vagin, V. V., Lee, S., Xu, J., Ma, S., Xi, H., Seitz, H., Horwich, M. D., Syrzycka, M., Honda, B. M., Kittler, E. L., Zapp, M. L., Klattenhoff, C., Schulz, N., Theurkauf, W. E., Weng, Z., and Zamore, P. D. (2009). Collapse of Germline piRNAs in the Absence of Argonaute3 Reveals Somatic piRNAs in Flies. Cell 137, 509–521.

Li, H. (2018). Minimap2: pairwise alignment for nucleotide sequences. Bioinformatics 34, 3094–3100.

Lin, M. H., Jensen, M. K., Elrod, N. D., Chu, H. F., Haseley, M., Beam, A. C., Huang, K. L., Chiang, W., Russell, W. K., Williams, K., Pröschel, C., Wagner, E. J., and Tong, L. (2024). Cytoplasmic binding partners of the Integrator endonuclease INTS11 and its paralog CPSF73 are required for their nuclear function. Mol Cell S1097-2765(24)00524.

Livneh, I., Cohen-Kaplan, V., Cohen-Rosenzweig, C., Avni, N., and Ciechanover, A. (2016). The life cycle of the 26S proteasome: from birth, through regulation and function, and onto its death. Cell Res 26, 869–885.

Lubas, M., Christensen, M. S., Kristiansen, M. S., Domanski, M., Falkenby, L. G., Lykke-Andersen, S., Andersen, J. S., Dziembowski, A., and Jensen, T. H. (2011). Interaction profiling identifies the human nuclear exosome targeting complex. Mol Cell 43, 624–637.

Lykke-Andersen, S., Žumer, K., Molska, E. Š., Rouvière, J. O., Wu, G., Demel, C., Schwalb, B., Schmid, M., Cramer, P., and Jensen, T. H. (2021). Integrator is a genome-wide attenuator of non-productive transcription. Mol Cell 81, 514–529.e6.

Matera, A. G., and Wang, Z. (2014). A day in the life of the spliceosome. Nat Rev Mol Cell Biol 15, 108–121.

Meola, N., Domanski, M., Karadoulama, E., Chen, Y., Gentil, C., Pultz, D., Vitting-Seerup, K., Lykke-Andersen, S., Andersen, J. S., Sandelin, A., and Jensen, T. H. (2016). Identification of a Nuclear Exosome Decay Pathway for Processed Transcripts. Mol Cell 64, 520–533.

Millevoi, S., and Vagner, S. (2010). Molecular mechanisms of eukaryotic pre-mRNA 3’ end processing regulation. Nucleic Acids Res 38, 2757–2774.

Mitchell, P., Petfalski, E., Shevchenko, A., Mann, M., and Tollervey, D. (1997). The exosome: a conserved eukaryotic RNA processing complex containing multiple 3’-->5’ exoribonucleases. Cell 91, 457–466.

Morawe, T., Honemann-Capito, M., von Stein, W., and Wodarz, A. (2011). Loss of the extraproteasomal ubiquitin receptor Rings lost impairs ring canal growth in *Drosophila* oogenesis. J Cell Biol 193, 71–80.

Morlan, J. D., Qu, K., and Sinicropi, D. V. (2012). Selective depletion of rRNA enables whole transcriptome profiling of archival fixed tissue. PLoS One 7, e42882.

Navarro-Costa, P., McCarthy, A., Prudêncio, P., Greer, C., Guilgur, L. G., Becker, J. D., Secombe, J., Rangan, P., and Martinho, R. G. (2016). Early programming of the oocyte epigenome temporally controls late prophase I transcription and chromatin remodelling. Nat Commun 7, 12331.

Neuman-Silberberg, F. S., and Schüpbach, T. (1993). The *Drosophila* dorsoventral patterning gene gurken produces a dorsally localized RNA and encodes a TGF alpha-like protein. Cell 75, 165–174.

Neuman-Silberberg, F. S., and Schupbach, T. (1994). Dorsoventral axis formation in *Drosophila* depends on the correct dosage of the gene gurken. Development 120, 2457–2463.

O’Reilly, D., Kuznetsova, O. V., Laitem, C., Zaborowska, J., Dienstbier, M., and Murphy, S. (2014). Human snRNA genes use polyadenylation factors to promote efficient transcription termination. Nucleic Acids Res 42, 264–275.

Offley, S. R., Pfleiderer, M. M., Zucco, A., Fraudeau, A., Welsh, S. A., Razew, M., Galej, W. P., and Gardini, A. (2023). A combinatorial approach to uncover an additional Integrator subunit. Cell Rep 42, 112244.

Pellizzoni, L., Yong, J., and Dreyfuss, G. (2002). Essential role for the SMN complex in the specificity of snRNP assembly. Science 298, 1775–1779.

Pfleiderer, M. M., and Galej, W. P. (2021). Structure of the catalytic core of the Integrator complex. Mol Cell 81, 1246–1259.e8.

Polák, P., Garland, W., Rathore, O., Schmid, M., Salerno-Kochan, A., Jakobsen, L., Gockert, M., Gerlach, P., Silla, T., Andersen, J. S., Conti, E., and Jensen, T. H. (2023). Dual agonistic and antagonistic roles of ZC3H18 provide for co-activation of distinct nuclear RNA decay pathways. Cell Rep 42, 113325.

Port, F., Chen, H.-M., Lee, T., and Bullock, S. L. (2014). Optimized CRISPR/Cas tools for efficient germline and somatic genome engineering in *Drosophila*. Proc Natl Acad Sci U S A 111, E2967–76.

Puno, M. R., and Lima, C. D. (2018). Structural basis for MTR4-ZCCHC8 interactions that stimulate the MTR4 helicase in the nuclear exosome-targeting complex. Proc Natl Acad Sci U S A 115, E5506–E5515.

Puno, M. R., and Lima, C. D. (2022). Structural basis for RNA surveillance by the human nuclear exosome targeting (NEXT) complex. Cell 185, 2132–2147.e26.

Quinlan, A. R., and Hall, I. M. (2010). BEDTools: a flexible suite of utilities for comparing genomic features. Bioinformatics 26, 841–842.

Ramírez, F., Ryan, D. P., Grüning, B., Bhardwaj, V., Kilpert, F., Richter, A. S., Heyne, S., Dündar, F., and Manke, T. (2016). deepTools2: a next generation web server for deep-sequencing data analysis. Nucleic Acids Res 44, W160–5.

Reits, E. A., Benham, A. M., Plougastel, B., Neefjes, J., and Trowsdale, J. (1997). Dynamics of proteasome distribution in living cells. EMBO J 16, 6087–6094.

Rennie, S., Dalby, M., Lloret-Llinares, M., Bakoulis, S., Dalager Vaagensø, C., Heick Jensen, T., and Andersson, R. (2018). Transcription start site analysis reveals widespread divergent transcription in *D. melanogaster* and core promoter-encoded enhancer activities. Nucleic Acids Res 46, 5455–5469.

Robinson, J. T., Thorvaldsdottir, H., Winckler, W., Guttman, M., Lander, E. S., Getz, G., and Mesirov, J. P. (2011). Integrative genomics viewer. Nat Biotechnol 29, 24–26.

Rosa-Mercado, N. A., Zimmer, J. T., Apostolidi, M., Rinehart, J., Simon, M. D., and Steitz, J. A. (2021). Hyperosmotic stress alters the RNA polymerase II interactome and induces readthrough transcription despite widespread transcriptional repression. Mol Cell 81, 502–513.e4.

Rubtsova, M. P., Vasilkova, D. P., Moshareva, M. A., Malyavko, A. N., Meerson, M. B., Zatsepin, T. S., Naraykina, Y. V., Beletsky, A. V., Ravin, N. V., and Dontsova, O. A. (2019). Integrator is a key component of human telomerase RNA biogenesis. Sci Rep 9, 1701.

Sabath, K., Qiu, C., and Jonas, S. (2024). Assembly mechanism of Integrator’s RNA cleavage module. Mol Cell S1097-2765(24)00539.

Sabath, K., Stäubli, M. L., Marti, S., Leitner, A., Moes, M., and Jonas, S. (2020). INTS10-INTS13-INTS14 form a functional module of Integrator that binds nucleic acids and the cleavage module. Nat Commun 11, 3422.

Sabin, L. R., Zhou, R., Gruber, J. J., Lukinova, N., Bambina, S., Berman, A., Lau, C. K., Thompson, C. B., and Cherry, S. (2009). Ars2 regulates both miRNA- and siRNA-dependent silencing and suppresses RNA virus infection in *Drosophila*. Cell 138, 340–351.

Saffman, E. E., Styhler, S., Rother, K., Li, W., Richard, S., and Lasko, P. (1998). Premature translation of oskar in oocytes lacking the RNA-binding protein bicaudal-C. Mol Cell Biol 18, 4855–4862.

Sánchez-Escabias, E., Guerrero-Martínez, J. A., and Reyes, J. C. (2022). Co-transcriptional splicing efficiency is a gene-specific feature that can be regulated by TGFβ. Commun Biol 5, 277.

Saunders, C., and Cohen, R. S. (1999). The role of oocyte transcription, the 5’UTR, and translation repression and derepression in *Drosophila* gurken mRNA and protein localization. Mol Cell 3, 43–54.

Schilders, G., Raijmakers, R., Raats, J. M. H., and Pruijn, G. J. M. (2005). MPP6 is an exosome-associated RNA-binding protein involved in 5.8S rRNA maturation. Nucleic Acids Res 33, 6795–6804.

Schmidt, D., Reuter, H., Hüttner, K., Ruhe, L., Rabert, F., Seebeck, F., Irimia, M., Solana, J., and Bartscherer, K. (2018). The Integrator complex regulates differential snRNA processing and fate of adult stem cells in the highly regenerative planarian *Schmidtea mediterranea*. PLoS Genet 14, e1007828.

Schulze, W. M., and Cusack, S. (2017). Structural basis for mutually exclusive co-transcriptional nuclear cap-binding complexes with either NELF-E or ARS2. Nat Commun 8, 1302.

Schwämmle, V., Hagensen, C. E., Rogowska-Wrzesinska, A., and Jensen, O. N. (2020). PolySTest: Robust Statistical Testing of Proteomics Data with Missing Values Improves Detection of Biologically Relevant Features. Mol Cell Proteomics 19, 1396–1408.

Sigova, A. A., Mullen, A. C., Molinie, B., Gupta, S., Orlando, D. A., Guenther, M. G., Almada, A. E., Lin, C., Sharp, P. A., Giallourakis, C. C., and Young, R. A. (2013). Divergent transcription of long noncoding RNA/mRNA gene pairs in embryonic stem cells. Proc Natl Acad Sci U S A 110, 2876–2881.

Sinsimer, K. S., Jain, R. A., Chatterjee, S., and Gavis, E. R. (2011). A late phase of germ plasm accumulation during *Drosophila* oogenesis requires lost and rumpelstiltskin. Development 138, 3431–3440.

Skaar, J. R., Ferris, A. L., Wu, X., Saraf, A., Khanna, K. K., Florens, L., Washburn, M. P., Hughes, S. H., and Pagano, M. (2015). The Integrator complex controls the termination of transcription at diverse classes of gene targets. Cell Res 25, 288–305.

Sun, M., Larivière, L., Dengl, S., Mayer, A., and Cramer, P. (2010). A tandem SH2 domain in transcription elongation factor Spt6 binds the phosphorylated RNA polymerase II C-terminal repeat domain (CTD). J Biol Chem 285, 41597–41603.

Szczepińska, T., Kalisiak, K., Tomecki, R., Labno, A., Borowski, L. S., Kulinski, T. M., Adamska, D., Kosinska, J., and Dziembowski, A. (2015). DIS3 shapes the RNA polymerase II transcriptome in humans by degrading a variety of unwanted transcripts. Genome Res 25, 1622–1633.

Tarn, W. Y., Hsu, C. H., Huang, K. T., Chen, H. R., Kao, H. Y., Lee, K. R., and Cheng, S. C. (1994). Functional association of essential splicing factor(s) with PRP19 in a protein complex. EMBO J 13, 2421–2431.

Tatomer, D. C., Elrod, N. D., Liang, D., Xiao, M. S., Jiang, J. Z., Jonathan, M., Huang, K. L., Wagner, E. J., Cherry, S., and Wilusz, J. E. (2019). The Integrator complex cleaves nascent mRNAs to attenuate transcription. Genes Dev 33, 1525–1538.

Tiedje, C., Lubas, M., Tehrani, M., Menon, M. B., Ronkina, N., Rousseau, S., Cohen, P., Kotlyarov, A., and Gaestel, M. (2015). p38MAPK/MK2-mediated phosphorylation of RBM7 regulates the human nuclear exosome targeting complex. RNA 21, 262–278.

Tilley, F. C., Arrondel, C., Chhuon, C., Boisson, M., Cagnard, N., Parisot, M., Menara, G., Lefort, N., Guerrera, I. C., Bole-Feysot, C., Benmerah, A., Antignac, C., and Mollet, G. (2021). Disruption of pathways regulated by Integrator complex in Galloway-Mowat syndrome due to WDR73 mutations. Sci Rep 11, 5388.

Tomecki, R., Kristiansen, M. S., Lykke-Andersen, S., Chlebowski, A., Larsen, K. M., Szczesny, R. J., Drazkowska, K., Pastula, A., Andersen, J. S., Stepien, P. P., Dziembowski, A., and Jensen, T. H. (2010). The human core exosome interacts with differentially localized processive RNases: hDIS3 and hDIS3L. EMBO J 29, 2342–2357.

Tomko, R. J., and Hochstrasser, M. (2011). Order of the proteasomal ATPases and eukaryotic proteasome assembly. Cell Biochem Biophys 60, 13–20.

Uriarte, M., Sen Nkwe, N., Tremblay, R., Ahmed, O., Messmer, C., Mashtalir, N., Barbour, H., Masclef, L., Voide, M., Viallard, C., Daou, S., Abdelhadi, D., Ronato, D., Paydar, M., Darracq, A., Boulay, K., Desjardins-Lecavalier, N., Sapieha, P., Masson, J.-Y., Sergeev, M., Kwok, B. H., Hulea, L., Mallette, F. A., Milot, E., Larrivée, B., Wurtele, H., and Affar, E. B. (2021). Starvation-induced proteasome assemblies in the nucleus link amino acid supply to apoptosis. Nat Commun 12, 6984.

von Mikecz, A. (2006). The nuclear ubiquitin-proteasome system. J Cell Sci 119, 1977–1984.

Vos, S. M., Farnung, L., Boehning, M., Wigge, C., Linden, A., Urlaub, H., and Cramer, P. (2018). Structure of activated transcription complex Pol II-DSIF-PAF-SPT6. Nature 560, 607–612.

Wasmuth, E. V., Januszyk, K., and Lima, C. D. (2014). Structure of an Rrp6-RNA exosome complex bound to poly(A) RNA. Nature 511, 435–439.

Wick, R. R., Judd, L. M., Gorrie, C. L., and Holt, K. E. (2017). Completing bacterial genome assemblies with multiplex MinION sequencing. Microb Genom 3, e000132.

Wilson, M. D., Wang, D., Wagner, R., Breyssens, H., Gertsenstein, M., Lobe, C., Lu, X., Nagy, A., Burke, R. D., Koop, B. F., and Howard, P. L. (2008). ARS2 is a conserved eukaryotic gene essential for early mammalian development. Mol Cell Biol 28, 1503–1514.

Winczura, K., Schmid, M., Iasillo, C., Molloy, K. R., Harder, L. M., Andersen, J. S., LaCava, J., and Jensen, T. H. (2018). Characterizing ZC3H18, a Multi-domain Protein at the Interface of RNA Production and Destruction Decisions. Cell Rep 22, 44–58.

Wu, G., Schmid, M., Rib, L., Polak, P., Meola, N., Sandelin, A., and Jensen, T. H. (2020). A Two-Layered Targeting Mechanism Underlies Nuclear RNA Sorting by the Human Exosome. Cell Rep 30, 2387–2401.e5.

Wu, Y., Albrecht, T. R., Baillat, D., Wagner, E. J., and Tong, L. (2017). Molecular basis for the interaction between Integrator subunits IntS9 and IntS11 and its functional importance. Proc Natl Acad Sci U S A 114, 4394–4399.

Yamamoto, J., Hagiwara, Y., Chiba, K., Isobe, T., Narita, T., Handa, H., and Yamaguchi, Y. (2014). DSIF and NELF interact with Integrator to specify the correct post-transcriptional fate of snRNA genes. Nat Commun 5, 4263.

Yang, J., Li, J., Miao, L., Gao, X., Sun, W., Linghu, S., Ren, G., Peng, B., Chen, S., Liu, Z., Wang, B., Dong, A., Huang, D., Yuan, J., Dang, Y., and Lai, F. (2024). Transcription directionality is licensed by Integrator at active human promoters. Nat Struct Mol Biol 31, 1208–1221.

Zhang, Z., Theurkauf, W. E., Weng, Z., and Zamore, P. D. (2012). Strand-specific libraries for high throughput RNA sequencing (RNA-Seq) prepared without poly(A) selection. Silence 3, 9.

Zheng, H., Jin, Q., Wang, X., Qi, Y., Liu, W., Ren, Y., Zhao, D., Xavier Chen, F., Cheng, J., Chen, X., and Xu, Y. (2023). Structural basis of INTAC-regulated transcription. Protein Cell 14, 698–702.

Zheng, H., Qi, Y., Hu, S., Cao, X., Xu, C., Yin, Z., Chen, X., Li, Y., Liu, W., Li, J., Wang, J., Wei, G., Liang, K., Chen, F. X., and Xu, Y. (2020). Identification of Integrator-PP2A complex (INTAC), an RNA polymerase II phosphatase. Science 370, eabb5872.

Zinder, J. C., Wasmuth, E. V., and Lima, C. D. (2016). Nuclear RNA Exosome at 3.1 Å Reveals Substrate Specificities, RNA Paths, and Allosteric Inhibition of Rrp44/Dis3. Mol Cell 64, 734–745.

